# A DIA-based quantitative crosslinking mass spectrometry framework for dynamic structural proteomics

**DOI:** 10.64898/2026.06.16.732730

**Authors:** Micha J. Birklbauer, Sowmya Sivakumar Geetha, Paul Getreuer, Gerlinde Grabmann, David Hollenstein, Wolfgang Kandioller, Viktoria Dorfer, Verena Jantsch, Karl Mechtler, Fränze Müller

**Affiliations:** Bioinformatics Research Group, University of Applied Sciences Upper Austria, Softwarepark 11, Hagenberg, 4232, Austria; Max Perutz Labs, Vienna BioCenter (VBC), Doktor-Bohr Gasse 9, Vienna, 1030, Austria; University of Vienna, Universitätsring 1, Vienna, 1030, Austria; Vienna BioCenter PhD Program, a Doctoral School of the University of Vienna and the Medical University of Vienna, Vienna, 1030, Austria; Institute of Inorganic Chemistry, University of Vienna, Währinger Straße 42, Vienna, 1090, Austria; Metabolomics, Vienna BioCenter Core Facilities GmbH, Vienna, Austria; Mass Spectrometry Facility, Max Perutz Labs, Dr.-Bohr-Gasse 9, Vienna, 1030, Austria; Institute of Molecular Biotechnology (IMBA), Vienna BioCenter (VBC), Doktor-Bohr Gasse 3, Vienna, 1030, Austria; Gregor Mendel Institute of Molecular Plant Biology (GMI), Vienna BioCenter (VBC), Doktor-Bohr Gasse 3, Vienna, 1030, Austria; Research Institute of Molecular Pathology (IMP), Vienna BioCenter (VBC), Campus-Vienna-Biocenter 1, Vienna, 1030, Austria

**Keywords:** Quantitative crosslinking mass spectrometry (QCLMS), Data-independent acquisition (DIA), Structural proteomics, Spectral library, Protein–RNA interactions, Conformational dynamics

## Abstract

Proteins undergo dynamic conformational rearrangements and interactions that are central to their biological functions. Quantitative crosslinking mass spectrometry enables the analysis of those dynamics and molecular interactions, but rigorous confidence assessment and empirical validation strategies for quantitative measurements remain underdeveloped, and integrated analysis of complementary structural features, including monolinks and protein–RNA adducts, remains limited. Here we present a data-independent acquisition (DIA)-based framework for quantitative crosslinking mass spectrometry (DIA-QCLMS) that combines optimized acquisition strategies, crosslink-aware spectral libraries and empirical false-discovery-rate (FDR) validation. The workflow supports crosslinks, monolinks and protein–RNA adducts and integrates spectral-library generation from two crosslinking search engines (xiSEARCH and MS Annika). To enable robust confidence assessment in DIA data, we developed a four-state target–decoy spectral library strategy that explicitly models target–target, target–decoy, decoy–target and decoy–decoy crosslink spectra. Experimental entrapment datasets enabled empirical validation of confidence estimation, whereas benchmarking with PhoX-crosslinked Cas9 demonstrated improved quantitative completeness and reproducibility compared with data-dependent acquisition. Application of the workflow to the ATP-dependent RNA helicase UAP56 (DDX39B) resolved ligand-dependent changes in intramolecular restraints, residue accessibility and candidate RNA-contact sites associated with the transition from an open to a clamped conformation. These results establish DIA-QCLMS as a scalable framework for quantitative structural proteomics and provide practical strategies for confidence-controlled analysis of dynamic protein interactions and conformational states.

## 1 Introduction

Proteins are central to practically all cellular processes, acting as dynamic molecular machines that couple structural rearrangements to biological function[1, 2]. Understanding protein structure and dynamics is therefore essential for elucidating mechanisms underlying health and disease[3, 4]. Protein dynamics extend beyond conformational changes and include processes such as metabolite binding and allosteric regulation[5, 6], post-translational modifications (PTMs)[7], protein–protein interactions (PPIs)[8, 9], and interactions with nucleic acids[10, 11], collectively defining the functional plasticity of the proteome[12]. In vivo, proteins operate within dynamic assemblies that respond to cellular signals[12–14], and disruption of these processes contributes to diseases such as cancer and neurodegeneration[15, 16].

Despite major advances in structural biology, capturing protein dynamics in native environments remains challenging. Although X-ray crystallography and cryo-electron microscopy can resolve multiple conformational states at high resolution, they typically rely on ensemble averaging and remain limited in detecting transient or low-population intermediates in complex biological systems[17, 18]. Similarly, traditional biochemical approaches often fail to detect weak or short-lived interactions[19]. To overcome these limitations, structural proteomics approaches, including crosslinking mass spectrometry[20–24], hydrogen–deuterium exchange (HDX)[25, 26], and limited proteolysis[27], have emerged as powerful tools to probe protein structure and dynamics under near-native conditions. Crosslinking mass spectrometry, in particular, provides spatial restraints by covalently linking proximal amino acid residues, thereby enabling the characterization of protein conformations and interaction networks. The integration of quantitative dimensions into crosslinking mass spectrometry (QCLMS) further allows the comparison of structural states and the detection of conformational changes in response to ligands or environmental stimuli[28–31]. QCLMS workflows have enabled robust quantification already in the past, allowing the investigation of enzyme dynamics[32, 33], protein complex rearrangements[29, 34], and ligand-induced structural changes[31, 35]. Recent methodological developments have significantly expanded the capabilities of QCLMS. Data-independent acquisition (DIA) strategies have improved reproducibility and quantitative accuracy, enabling large-scale and label-free analyses of crosslinked peptides[31, 36–40]. In this context, photoactivatable crosslinkers such as SDA or LCSDA provide additional advantages by enabling precise temporal control of crosslinking through UV activation and by capturing transient interactions with high efficiency[41–43]. These reagents are particularly well suited to study dynamic processes that involve rapid conformational transitions or weak interactions[41, 44–46].

The application of photo-triggered QCLMS provides a powerful framework to investigate conformational dynamics of for example ATP-dependent RNA helicases such as UAP56. UAP56 (DDX39B) is a core component of the TREX complex and plays a central role in coupling mRNA processing to nuclear export, undergoing ATP- and RNA-dependent conformational changes that are essential for its function[47–51]. Structural studies have shown that UAP56 transitions between open, half-open, and clamped states, in which its RecA1 and RecA2 domains rearrange to capture RNA and coordinate ATP hydrolysis[49, 50]. While cryo-electron microscopy and crystallographic approaches have provided high-resolution snapshots of these states, the dynamic coupling between RNA binding and ATPase activity remains incompletely understood[52]. Quantitative crosslinking approaches have the potential to address these limitations by capturing changes in domain proximity and protein–RNA interactions in response to nucleotide and substrate binding, thereby providing insight into the mechanistic basis of UAP56 function within the mRNA export pathway.

Here we established a DIA-based quantitative crosslinking mass spectrometry framework for dynamic structural proteomics. We hypothesized that combining optimized DIA acquisition, crosslink-aware spectral libraries, empirically benchmarked FDR control and photoactivatable chemistries would enable reproducible quantification of crosslinked peptides across purified proteins and complex biological samples. We first benchmarked acquisition parameters and quantitative accuracy using a PhoX-crosslinked Cas9 standard, then evaluated spectral-library validation and empirical FDR behavior using entrapment datasets derived from crosslinked *C. elegans* nuclei. Finally, we applied the workflow to the ATP-dependent RNA helicase UAP56 to test whether state-resolved QCLMS can capture ligand-dependent changes in intramolecular restraints, monolink accessibility and candidate RNA-proximal residues. Together, these experiments define a scalable strategy for measuring protein structural dynamics by DIA-QCLMS.

## 2 Results

### 2.1 A DIA-QCLMS workflow for reproducible quantification of crosslinked peptides

Here we present an extension of our previously established DIA-based quantitative crosslinking mass spectrometry (DIA-QCLMS) workflow[37, 38], implementing a peptide-centric analysis strategy to enable reproducible quantification of crosslinked peptides across diverse experimental systems (Figure 1). The workflow is compatible with both label-free and isotope-labeled approaches and focuses on optimized DIA acquisition and a flexible computational analysis pipeline. Sample preparation and enrichment strategies for crosslinking experiments have been described in detail elsewhere[28] and were therefore not modified here. The workflow relies on data dependent acquisition prior to DIA analysis for spectral library generation.

**Figure 1:**
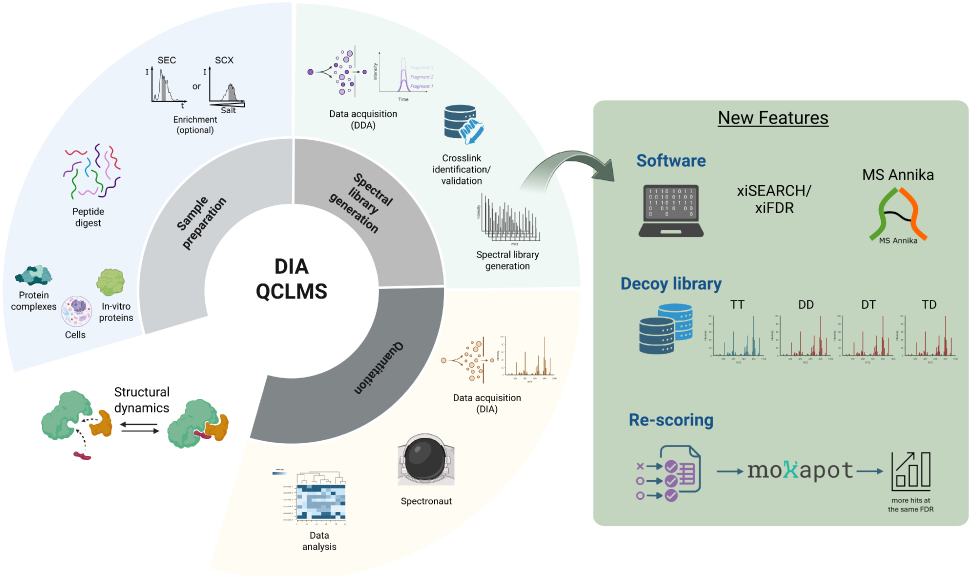
General Overview of the DIA-QCLMS workflow integrating sample preparation, spectral library generation and quantification. The left circle view was adapted from [38]. Created with BioRender.com.

To improve identification confidence and interoperability, the spectral library module was expanded to support multiple crosslink search platforms. In addition to xiSEARCH in combination with xiFDR, crosslinks can be identified using MS Annika within Proteome Discoverer, with FDR-controlled crosslink-spectrum matches (CSMs) serving as input for library generation. The library construction workflow was further extended to accept multiple input formats, including MGF and mzML files. Importantly, we implemented a crosslink-aware target–decoy strategy that generates target–target, target–decoy, decoy–target and decoy–decoy entries for each precursor, reflecting the paired nature of crosslinked peptides and enabling empirical assessment of false discovery rates in DIA analysis[53]. We further evaluated rescoring strategies for quantitative crosslinking data using mokapot, testing both peptide-level and crosslink-level feature sets derived from spectral library information. However, across the datasets examined, native Spectronaut validation provided the most consistent balance between identification confidence and quantitative performance, and was therefore used for subsequent analyses. Beyond intramolecular crosslinks, the workflow integrates monolinks and protein–RNA crosslinks to capture complementary aspects of protein structure and interaction. Monolinks provide proxies for local residue accessibility, whereas protein–RNA adducts report on RNA-proximal regions. To enable quantitative comparison of these features across conditions, we implemented a post-translational modification (PTM)-style analysis workflow in Spectronaut using the directDIA strategy[53, 54], allowing unified quantification of crosslinks, monolinks and protein–RNA adducts.

To evaluate the performance and general applicability of the workflow, we designed a three-tier experimental strategy spanning increasing levels of biological complexity (Figure 2).

**Figure 2:**
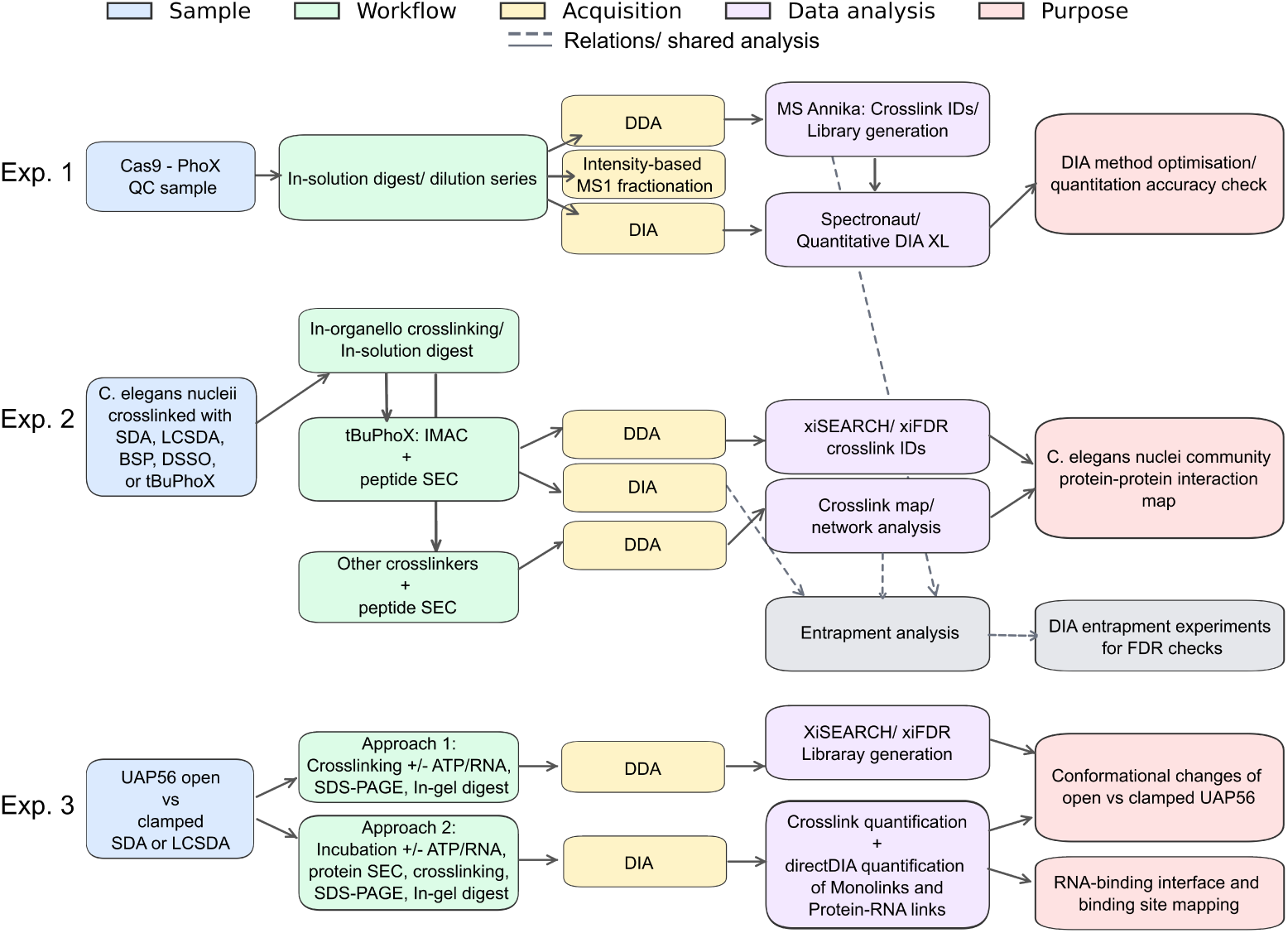
Experimental rationale overview. This study comprises three experimental tiers: first, DIA method development using a Cas9 crosslinked quality-control sample; second, generation of a community C. elegans nuclear protein–protein interaction map, which was additionally used for entrapment experiments and FDR assessment; and third, a biological showcase experiment integrating quantitative analysis of crosslinks, monolinks, and protein–RNA crosslinks.

In Experiment 1, we optimized DIA acquisition parameters using a single-protein quality control (QC) sample consisting of PhoX-crosslinked Cas9. To minimize variability arising from crosslinking, all injections were performed from a single preparation batch. A dilution series (1–500 ng) was analyzed to assess quantitative accuracy, chromatographic sampling density and reproducibility across technical replicates. Spectral libraries were generated from previously published DDA data[55] with the PRIDE identifier PXD059096, using custom scripts and multiple gradient lengths, with retention time alignment achieved via spiked-in iRT peptides[55, 56]. In Experiment 2, we applied the workflow to crosslinked nuclei isolated from *Caenorhabditis elegans* using multiple crosslinkers, including SDA, LCSDA, BSPNO2[57–59], also known as CLIP[57], DSSO, and tBuPhoX (commercially available as TBDSPP). Protein–protein interactions were captured by in-organelle crosslinking, followed by peptide fractionation (SDA, LCSDA, BSPNO2, and DSSO) or IMAC enrichment combined with fractionation for tBuPhoX. Crosslinks were identified and validated using xiSEARCH and xiFDR at 5% protein–protein interaction FDR. The resulting datasets were used to generate crosslinker-specific spectral libraries for entrapment analyses and to assemble a combined nuclear protein–protein interaction resource.

In Experiment 3, we applied the workflow to the ATP-dependent RNA helicase UAP56 (DDX39B), a DExD-box ATPase whose RecA1 and RecA2 domains adopt a clamped conformation on RNA in the presence of ATP[50]. UAP56 is proposed to function as a molecular bridge between the THO complex and messenger ribonucleoprotein particles (mRNPs), through connection of the RecA1 and RecA2 domains with the THOC2 subunit[49, 60, 61], and an additional interaction of the RecA domains to mRNPs via UAP56-binding-motifs found in export adapters such as ALYREF[62, 63]. To assess ligand-dependent structural changes, recombinant UAP56 was analyzed in an apo/open state and in the presence of ATP and poly(U) RNA, corresponding to a clamped conformation. Monolinks were used as complementary proxies for residue accessibility. Two complementary strategies were employed: (i) direct crosslinking of open and ligand-bound states using SDA or LCSDA, followed by SDS–PAGE separation and in-gel digestion; and (ii) incubation of UAP56 with ATP and 15U RNA followed by size-exclusion chromatography to isolate RNA-bound monomeric species prior to crosslinking. Spectral libraries were generated from DDA data and used for DIA-based quantification of crosslinks, protein–RNA adducts and monolinks.

### 2.2 A four-state target–decoy spectral library strategy for DIA-based crosslinking mass spectrometry

To enable confident identification and quantification of crosslinked peptides in DIA, we developed a crosslink-aware spectral library generation workflow that explicitly accounts for the paired nature of crosslinked peptides (Figure 3).

**Figure 3:**
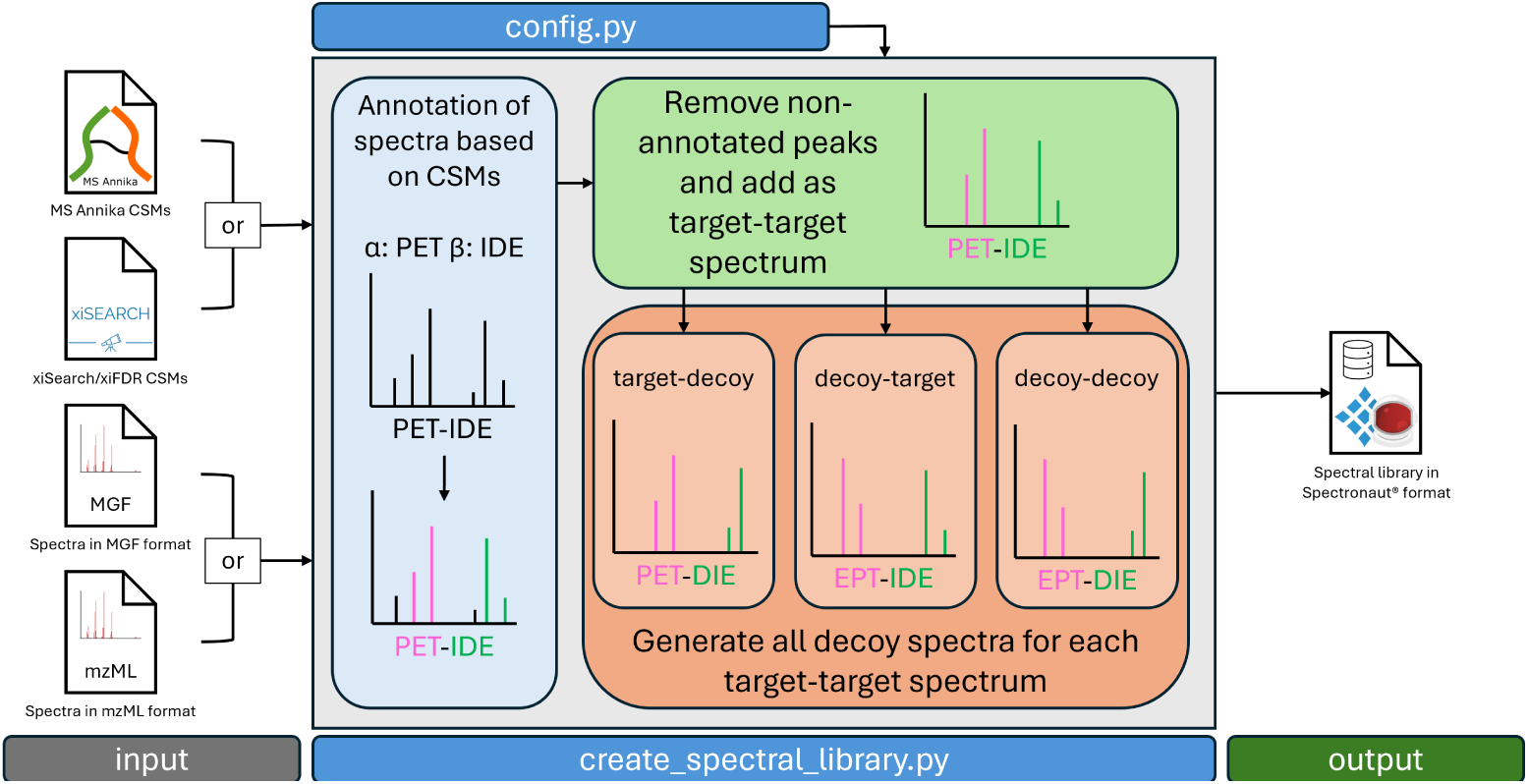
Schematic overview of the spectral library generation workflow. The whole workflow is implemented as a python script that takes crosslink-spectrum-matches (CSMs) from either MS Annika [64] or xiFDR [53] as input additionally to mass spectra in either MGF or mzML format and a configuration file. Mass spectra are annotated by calculating all theoretical fragment ions of the corresponding CSMs and any non-annotated peaks are removed before the spectrum is added to the spectral library. For every target spectrum we then calculate three different decoy spectra (target-decoy, decoy-target, and decoy-decoy) to capture the more complex behaviour of crosslinking validation. Decoys are generated using the reverse algorithm proposed by Zhang et al. [65], in short peptide sequences are reversed with exception of the C-terminal residue and peaks are shifted to their new positions. The final library containing target and decoy spectra (in proportion 1:3) is finally exported as a comma-separated-values file in the Spectronaut spectral library format.

The approach builds on data-dependent acquisition (DDA)-derived crosslink identifications and is designed for direct integration with Spectronaut for downstream DIA analysis. Spectral library construction proceeds in two steps. First, high-confidence crosslink-spectrum matches (CSMs) are used to generate annotated spectra by calculating all theoretical fragment ion masses for the identified peptide pair and matching them to experimental peaks. Only annotated fragment ions are retained, and non-assigned peaks are removed, yielding cleaned, crosslink-specific spectra that are incorporated as target entries in the spectral library. Second, to enable robust validation and false discovery rate (FDR) estimation, we implemented a four-state target–decoy strategy that reflects the unique properties of crosslinked peptide identification. In contrast to linear peptide proteomics, where each match is classified as either target or decoy, crosslinked peptide pairs can give rise to four distinct combinations: target–target (TT), target–decoy (TD), decoy–target (DT) and decoy–decoy (DD) matches[53]. To preserve this relationship in the DIA context, we added the three corresponding decoy spectra for each target spectrum to the spectral library.

Decoy spectra were created using a sequence-reversal approach as described by Zhang et al.[65], in which peptide sequences are reversed while retaining the C-terminal residue and preserving modifications on their specific amino acids. Fragment ion m/z values are recalculated according to the reversed sequences while conserving the original fragment ion intensities. For TD and DT spectra, this transformation is applied to only one of the two peptides, whereas DD spectra are generated by reversing both peptides. This results in a balanced spectral library containing TT, TD, DT and DD entries for each precursor, enabling empirical assessment of identification confidence in DIA-based crosslinking workflows. The resulting spectral libraries are exported in a comma-separated values (csv) format compatible with Spectronaut. The entire workflow is implemented as an open-source Python script with a command-line interface and supports multiple input formats, including CSM tables from MS Annika[64] or xiFDR[53], as well as mzML or MGF spectral data. The tool is freely available at https://github.com/hgb-bin-proteomics/MSAnnika_Spectral_Library_exporter. This strategy enables balanced representation of all target–decoy combinations and provides the basis for accurate empirical FDR estimation in our DIA-based crosslinking experiments.

### 2.3 DIA acquisition optimization balances crosslink coverage and quantitative precision

To establish a robust DIA acquisition strategy for quantitative crosslinking mass spectrometry, we systematically evaluated key parameters using a PhoX-crosslinked Cas9 quality-control sample (Figure 4). The performance was assessed based on three metrics relevant for quantitative analysis: the number of unique residue pairs (URPs), the number of data points sampled across chromatographic peaks, and the coefficient of variation (CV) between technical replicates. We first investigated the effect of DIA isolation window width on crosslink identification and sampling performance (Figure 4A–C). Narrow windows maximized crosslink identifications, with 5 m/z windows yielding 428 URPs on average, followed by 8 m/z (398 URPs), whereas broader windows progressively reduced identifications to 116 URPs at 20 m/z. This decrease is consistent with increased spectral complexity and reduced precursor selectivity at wider isolation windows. In contrast, wider windows improved chromatographic sampling, increasing the number of data points per peak from 3 (5 m/z) to 10.5 (20 m/z), corresponding to shorter duty cycles (0.78 s for 20 m/z versus 3.86 s for 5 m/z).

**Figure 4:**
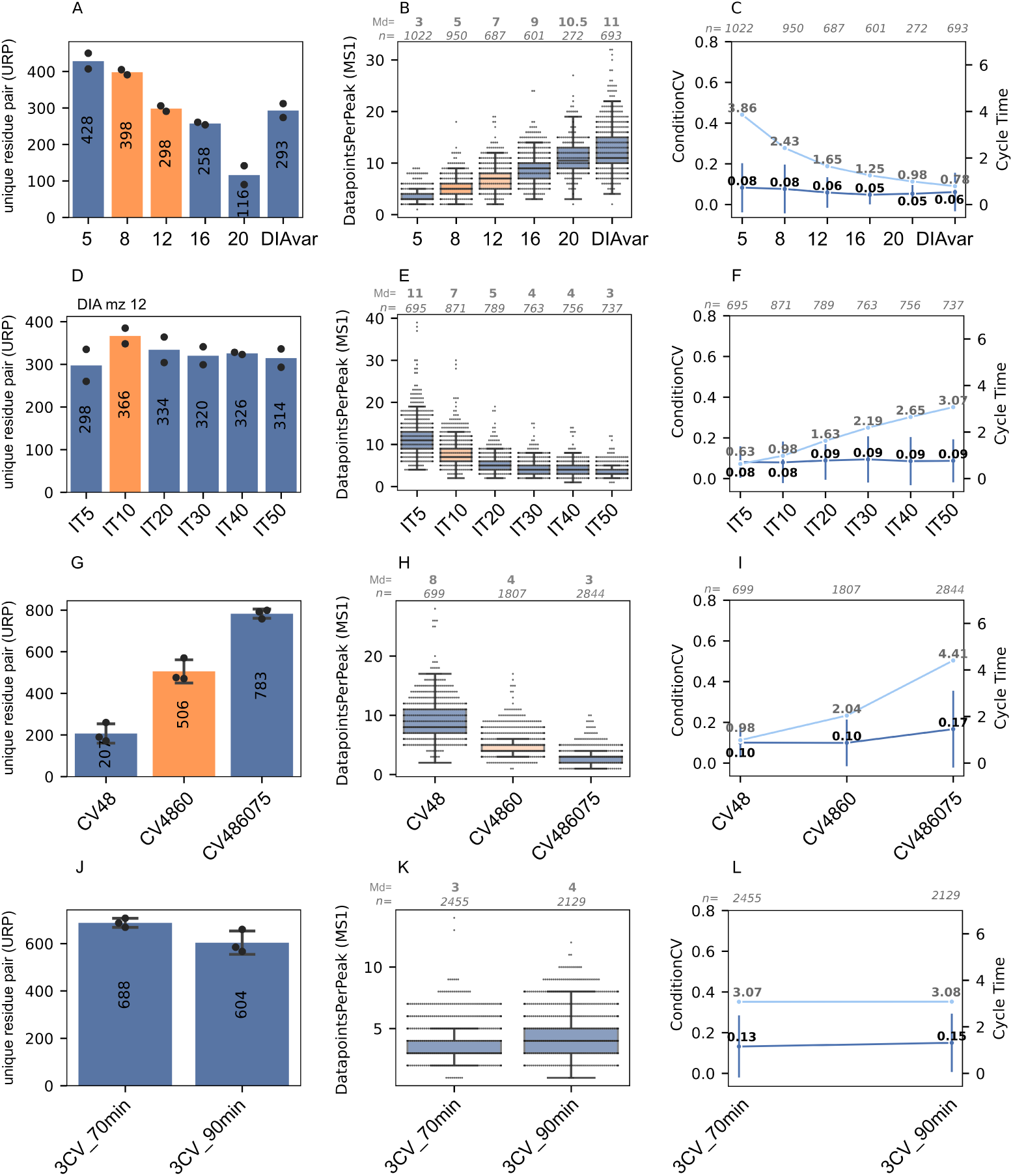
DIA method development for quantitative crosslinking mass spectrometry using 250 ng PhoX-crosslinked Cas9 QC samples. Systematic optimization of DIA acquisition parameters evaluating crosslink identifications and quantitative sampling performance across MS_1_ peaks. **A–C**, Effect of DIA window size (m/z 5, 8, 12, 16, 20, and variable windows) on unique residue pair (URP) identifications (**A**), data points per MS_1_ peak (**B**), and cycle time (**C**). Subsequent optimizations were performed using three methods: m/z 12, m/z 8 (Supplementary Fig. S1), and variable-window DIA (Supplementary Fig. S2). **D–F**, MS_2_ injection time optimization (5, 10, 20, 30, 40, and 50 ms), showing URP identifications (**D**), data points per MS_1_ peak (**E**), and cycle time (**F**). **G–I**, Compensation voltage (CV) optimization using FAIMS with a single CV (−48 V), double CVs (−48, −60 V), and triple CVs (−48, −60, −75 V). URP identifications (**G**), data points per MS_1_ peak (**H**), and coefficient of variation (CV) of quantified crosslinks (**I**) are shown. The CV combination was previously optimized[**?**]. **J–L**, Effect of gradient length (70 min versus 90 min) using three CVs, a fixed DIA window size (m/z 12), and a 10 ms MS_2_ injection time. URP identifications (**J**), data points per MS_1_ peak (**K**), and coefficient of variation (**L**) are shown. Measurements were performed in technical duplicates (**A–F**) or technical triplicates (**G–L**). For all bar plots, the mean number of unique residue pairs is indicated within each bar. Individual technical replicates are shown as black dots. Error bars (triplicates) represent the standard deviation, calculated as the average deviation of individual data points from the mean. Data-point distributions per MS_1_ peak were visualized using Seaborn boxplots with overlaid individual observations. In each boxplot, the center line represents the median (Md), the box boundaries indicate the interquartile range (IQR; 25th to 75th percentile), and whiskers extend to 1.5× the IQR. Individual data points are shown as jittered dots to illustrate distribution density and variability. For line plots, points with values represent the mean of technical replicates (n = 2 for **A–F**; n = 3 for **G–L**). Error bars indicate standard deviation (s.d.). No formal statistical hypothesis testing was performed; plots are shown for descriptive comparison. Numbers of individual observations are plotted above each boxplot and line plot.

These results highlight a fundamental trade-off between identification depth and quantitative sampling density. Although 5 m/z windows yielded the highest number of identifications, the limited number of data points per peak compromised peak definition and quantitative robustness. In contrast, 8 and 12 m/z window strategies maintained high crosslink coverage while providing sufficient sampling density (median 5–7 data points per peak) for reliable quantification. Based on this balance, subsequent optimization focused on these window sizes. We next evaluated the effect of MS2 injection time on identification and sampling performance (Figure 4D–F; Supplementary Figure S1). Intermediate injection times provided the best overall performance, with a maximum of 366 URPs observed at 10 ms. Increasing injection time reduced identification rates, likely due to longer cycle times and decreased chromatographic sampling frequency, whereas shorter injection times improved sampling density but did not further increase identification numbers. Accordingly, a 10 ms MS2 injection time was selected as a compromise between sensitivity and temporal resolution.

We then assessed the impact of FAIMS compensation voltage (CV) stepping on crosslink coverage (Figure 4G–I). Consistent with previous observations[55], combining multiple CV settings substantially increased crosslink identifications, with triple CV acquisition (−48, −60 and −75 V) yielding 783 URPs compared to 204 URPs for a single CV (−48 V). However, this gain in coverage was accompanied by reduced chromatographic sampling, as cycle times increased and the number of data points per peak decreased from approximately 8 to 3. This illustrates a second key trade-off between precursor coverage and quantitative precision. Finally, we evaluated the effect of chromatographic gradient length using the optimized three-CV method (Figure 4J–L). Extending the gradient from 70 to 90 min resulted in only minor differences in crosslink identifications (688 versus 604 URPs) and did not substantially affect sampling density, indicating that shorter gradients already provided sufficient separation under the tested conditions. Comparable trends were observed for 8 m/z isolation windows (Supplementary Figure S1D–F). Although dual and triple CV acquisition increased crosslink identifications, the resulting reduction in data points per peak limited quantitative performance. Consequently, the 8 m/z method was not pursued further.

We also explored a variable-window DIA strategy to further improve chromatographic sampling (Figure 4A; Supplementary Figure S2). This approach achieved similar crosslink identification rates compared to fixed 12 m/z windows while increasing the number of data points per peak from approximately 7 to 11. Additional optimization of MS1 injection time, MS2 injection time and automatic gain control (AGC) confirmed that cycle time was largely governed by the cumulative MS2 acquisition, consistent with previous observations on the Orbitrap Astral platform[66]. While shorter MS2 injection times further increased sampling density (up to 17 data points per peak), this did not translate into consistent improvements in crosslink identification. Despite the superior sampling performance of variable-window DIA, this approach required broad isolation windows at the edges of the m/z range, reducing precursor selectivity. We therefore did not apply this strategy for subsequent biological analyses and instead selected a fixed-window method to maintain consistent identification confidence across the full m/z range. Across all acquisition strategies, quantitative reproducibility remained high, with coefficients of variation generally below 10%. However, variability increased with the use of multiple CV settings, reaching up to 17%. Considering both identification coverage and quantitative precision, we selected a DIA method combining 12 m/z isolation windows, a 10 ms MS2 injection time and dual CV acquisition (−48/−60 V) as a balanced configuration for downstream analyses. This method resulted in 506 URPs, 4 data points per peak and 10% CV.

### 2.4 DIA improves quantitative completeness and reproducibility across crosslink abundance ranges

Additionally, we assessed the quantitative accuracy of DIA measurements using a dilution series of the PhoX-crosslinked Cas9 quality-control sample (Figure 5A-D). The sample was injected in triplicate at 1, 10, 50, 100, 250 and 500 ng. As expected, the 1 ng injection yielded the lowest number of crosslink identifications, with 18 unique residue pairs, and showed the highest variability between technical replicates (CV 18%). This resulted in the poorest quantitative accuracy, with a median log10 MS1 peak area ratio of −2.05 compared with the expected ratio (theoretical ratio to reference channel 500 ng) of −2.7. Increasing the injection amount to 10 ng improved performance to 34 URPs, with technical variability reduced to 5% and a measured MS1 peak area ratio of −1.67, closely matching the expected ratio of −1.7. With further increases in sample load, the number of identifications rose from 138 URPs at 50 ng to a maximum of 265 URPs at 500 ng. Technical variability remained low across higher injection amounts, ranging from 8% at 50 ng to 4% at 500 ng. This was also reflected in the quantitative accuracy, where median MS1 peak area ratios closely matched the respective expected ratios indicated by the red reference lines (Figure 5C). The number of data points per peak ranged from 7 to 8 across all injection amounts and remained stable throughout the series. From this experiment, we conclude that variability between technical replicates is closely linked to quantitative accuracy and therefore we minimized it throughout the experiment.

**Figure 5:**
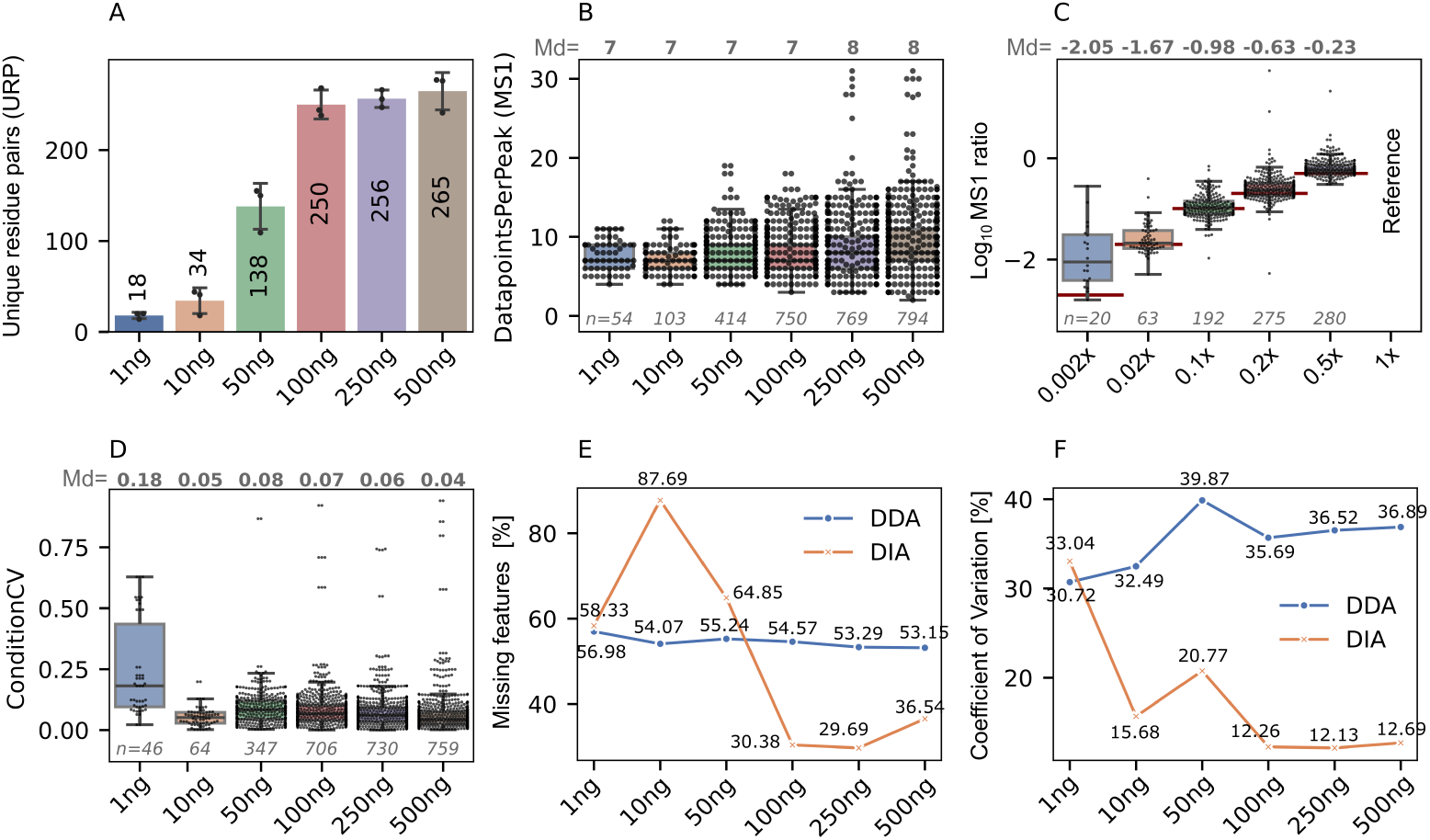
Accuracy of DIA-based quantitative crosslinking across a Cas9–PhoX dilution series. Quantitative evaluation of data-independent acquisition (DIA) crosslinking measurements of PhoX-crosslinked Cas9 across injection amounts ranging from 1–500 ng. **A**, Number of unique residue pairs (URPs) identified in DIA mode as a function of injection amount. Measurements were performed on an Orbitrap Astral mass spectrometer coupled to a Vanquish Neo system equipped with an Aurora Ultimate 25 cm column. **B**, Number of data points acquired per MS_1_ peak for each injection amount, showing consistently reproducible sampling with average values between 7.3 and 8.9 data points per peak. **C**, log_10_-transformed DIA MS_1_ peak area ratios across the dilution series (0.002× (1 ng) to 0.5× (250 ng)). Red lines indicate the expected theoretical ratios for each dilution step. **D**, Coefficient of variation (CV) of quantified crosslinks at each injection amount, demonstrating consistently low variability across the 10–500 ng range. **E**, Percentage of missing features across the dilution series. Features were considered missing if detected and quantified in fewer than three out of three technical replicates. DIA data are shown in orange and data-dependent acquisition (DDA) data in blue. **F**, Comparison of CV values across the dilution series between DIA (orange) and DDA (blue) crosslinking datasets, illustrating improved quantitative precision for DIA measurements. For the bar plot, the mean number of unique residue pairs is indicated within each bar. Individual technical replicates are shown as black dots. Error bars (triplicates) represent the standard deviation, calculated as the average deviation of individual data points from the mean. Data-point distributions per MS_1_ peak were visualized using Seaborn boxplots with overlaid individual observations. In each boxplot, the center line represents the median (Md), the box boundaries indicate the interquartile range (IQR; 25th to 75th percentile), and whiskers extend to 1.5× the IQR. Individual data points are shown as jittered dots to illustrate distribution density and variability. For line plots, points represent the mean of technical replicates (n = 3). For **E** and **F**, dots with values represent the mean value of a triplicate analysis (**F**) or the overall percentage of missing features (**E**).

We were also interested in comparing DDA and DIA crosslink measurements with respect to missing features (missing values) and technical variability across triplicate injections. To assess the proportion of missing values, triplicate datasets were filtered according to the number of observed replicates. If one value was missing within a triplicate set, the corresponding feature was classified as incomplete and counted as a missing feature. For each injection amount, the percentage of missing features was then calculated, reflecting the proportion of triplicates lacking complete quantitative information (Figure 5E). For DDA analysis, the proportion of missing features remained consistently high across all injection amounts, ranging from 53% (500 ng) to 57% (1 ng). This is attributable to the stochastic precursor selection inherent to DDA acquisition, in which peptide ions compete for MS/MS sampling in each cycle, resulting in incomplete and less reproducible feature sampling between replicate runs. This phenomenon has been widely reported in shotgun proteomics studies[67]. Although this issue can be overcome by propagating IDs between runs (comparable to match between runs in DIA) as described by Kalxdorf et al.[68], crosslink search engines have not integrated this mapping approach to this date. In contrast, DIA injections showed a high proportion of missing features at low injection amounts (1 ng, 58%; 10 ng, 88%; 50 ng, 65%), whereas missing values decreased markedly at higher injection amounts (100 ng, 30%; 250 ng, 30%; 500 ng, 37%) probably due to the superior performance of match between run across technical replicates. Thus, DIA measurements became substantially more complete at injection amounts of 100 ng and above. A similar trend was observed for the coefficients of variation calculated as means across triplicate injections and residue pairs. In DDA measurements, variability increased at higher injection amounts from 31% (1 ng) to 37% (500 ng), with a maximum of 40% at 50 ng. In contrast, mean variability in DIA measurements decreased with increasing injection amount, from 33% (1 ng) to 13% (500 ng). Notably, already at an injection amount of 10 ng, DIA measurements were more reproducible than DDA acquisitions. Although the overall number of crosslink identifications obtained by DIA was lower than that achieved by DDA[55], missing features were substantially reduced at higher injection amounts and technical variability was lower from 10 ng onwards. We therefore conclude that DIA-based quantification of crosslinks improves measurement completeness, quantitative accuracy, and reproducibility. This is particularly advantageous for assessing differential crosslink abundances to resolve conformational changes and structural rearrangements.

We also adapted our workflow to Skyline, an open source software for target peptide analysis[69, 70]. Our integration is based on previous work using Skyline for crosslink quantitation[71] and recent implementation of DIA crosslinking data acquired on a TimsTOF device using Skyline for quantitation[39]. For spectral-library-based analysis, we generated a custom .ssl library and imported it into Skyline, followed by DIA raw data for automated extraction and quantification of crosslinked peptides. As Skyline currently does not support decoy spectral libraries for this workflow, data were filtered using idotp (isotope dot product, spectral similarity between areas of precursor isotopic peaks and expected isotopic profile) thresholds of 0.7 and 0.95 to control confidence. 0.95 indicates a high degree of similarity, providing confidence that the correct peptide is being identified and quantified. We also selected 0.7 as a relaxed quality threshold to compare the effect of data confidence. The number of identified unique peptide pairs increased with injection amount and plateaued at 100 ng (Supplementary Figure S4A,D), with up to 616 unique peptide pairs (UPP) at idotp 0.7 and 356 UPP at idotp 0.95, indicating consistent sensitivity scaling. Quantitative accuracy, assessed by median *log*_10_ MS1 peak area ratios, followed expected trends across the dilution series, with improved agreement at higher inputs but higher variability at lower stringency (idotp 0.7) (Supplementary Figure S4B, E). Remarkably, peak area ratios at dilution of 0.002x matched the expected ratios more closely as compared to our Spectronaut analysis. Variability was reduced using stricter filtering (idotp 0.95), although overall coefficients of variation remained elevated (Supplementary Figure S4C, F). We see strong potential for this workflow in DIA-based crosslink quantification and propose that incorporating decoy spectral libraries into Skyline-based analyses could further improve quantitative accuracy and reduce variability, enabling more robust high-throughput DIA crosslink analysis without manual peak curation.

### 2.5 Multi-chemistry crosslinking of *C. elegans* nuclei provides a complex benchmark and reusable interaction resource

Accurate false discovery rate (FDR) control is essential in crosslinking mass spectrometry because large peptide-pair search spaces increase the risk of false-positive identifications[53]. Most workflows estimate FDR using target–decoy strategies, in which spectra are searched against target and artificial decoy peptide pairs. Reported FDR reflects the expected proportion of false positives among accepted crosslink identifications. Incorrect FDR control can lead to spurious residue pairs or inflated protein interaction networks and may bias comparisons between methods[72]. To independently evaluate FDR performance, entrapment strategies can be used by adding peptides or proteins known to be absent from the sample[73, 74] to the search. Identifications assigned to this entrapment set represent false positives and provide an empirical benchmark for validating target–decoy-based FDR estimation. To evaluate FDR control of DIA crosslinking data and our decoy spectral library approach we generated three crosslinking spectral library data sets for entrapment experiments (i) *C. elegans* nuclei crosslinked with either, SDA, LCSDA, BSPNO2, DSSO or tBuPhoX, (ii) Human ribosomes and *E. coli* ribosomes crosslinked with PhoX and (iii) *C. elegans* nuclei crosslinked with DSG. The crosslinked *C. elegans* nuclei also serve as a community resource for the *C. elegans* community to retrieve protein-protein interactions (PPIs) for biological studies.

To generate the first entrapment dataset, *C. elegans* nuclei were isolated from worms and crosslinked using five different crosslinkers. Following nuclear crosslinking, samples were digested in solution, yielding soluble and insoluble digestion fractions. The soluble fraction was subjected to peptide size-exclusion chromatography (SEC) for enrichment of crosslinked peptides. In the case of tBuPhoX, crosslinked peptides were first enriched by immobilized metal affinity chromatography (IMAC) prior to SEC. The insoluble fraction was further processed using a filter-aided sample preparation (FASP) protocol to assess crosslinked proteins requiring stronger solubilization before enzymatic digestion. This fraction was subsequently also subjected to SEC for additional crosslinked peptide enrichment. To generate experimental entrapment spectral libraries, samples were analyzed by data-dependent acquisition, whereas only the tBuPhoX nuclear crosslinking samples were additionally measured by DIA. The DDA measurements yielded 578 and 733 URPs for the tBuPhoX and BSPNO2 entrapment datasets, respectively (Figure 6A).

**Figure 6:**
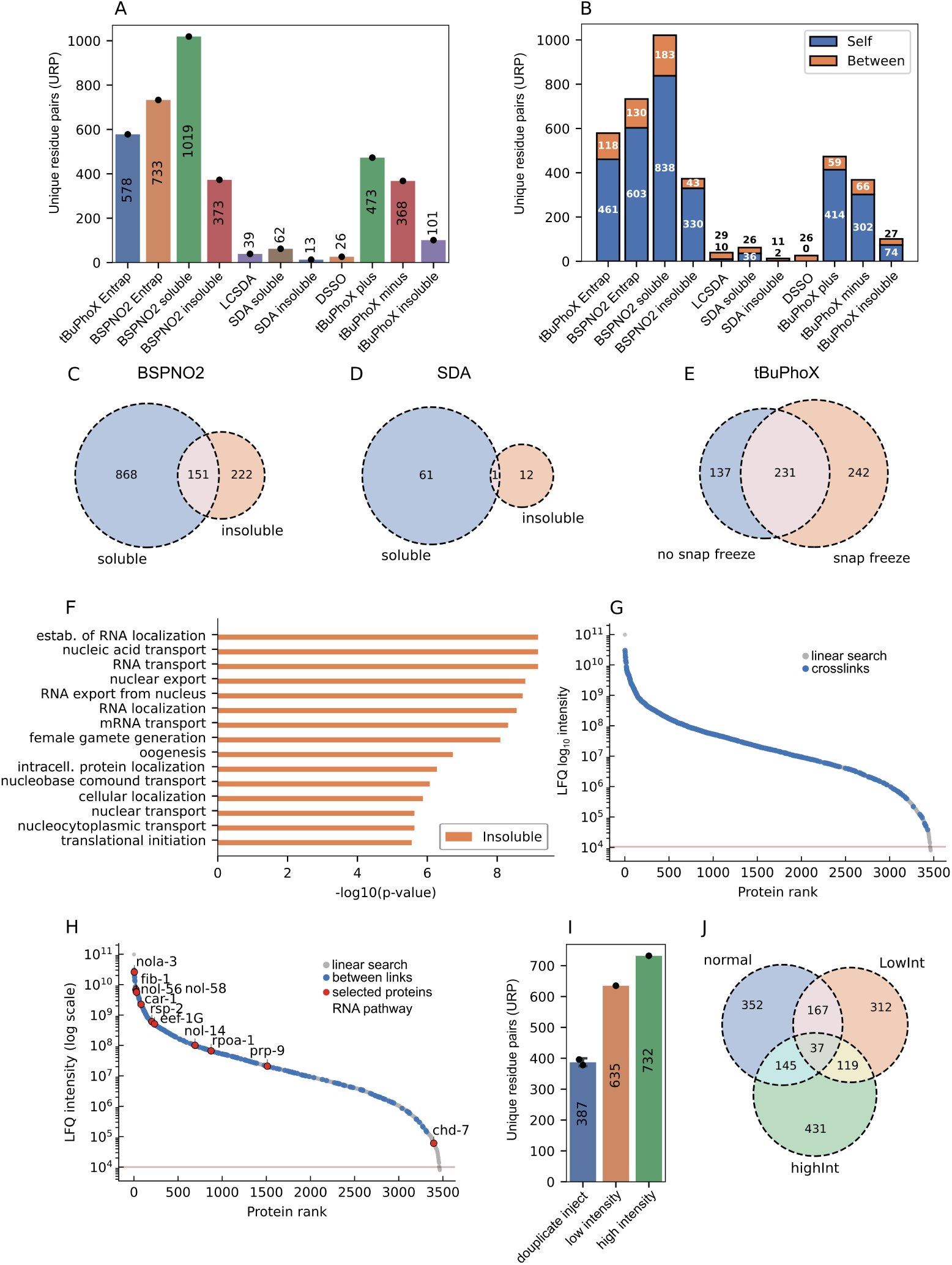
C. elegans nuclei crosslinking experiments using five different crosslinkers (tBuPhoX, BSPNO2, SDA, LCSDA, and DSSO) for entrapment analyses and generation of a community C. elegans protein–protein interaction network resource. **A**, Number of unique residue pairs (URPs) identified in individual C. elegans nuclei crosslinking experiments using different crosslinkers and peptide-fractionation strategies. Bars represent mean values from SEC-fractionation measurements. **B**, Distribution of intraprotein (“Self” or intra, blue) and interprotein (“Between” or inter, orange) crosslinks for the corresponding datasets shown in **A**. Numbers within bar segments indicate counts of each crosslink class. For the LCSDA, SDA insoluble, and DSSO datasets, values are plotted outside the bars because of low counts; the upper number indicates interprotein links and the lower number intraprotein links. **C–E**, Pairwise overlap of URPs between BSPNO2 soluble and insoluble fractions (**C**), SDA soluble and insoluble fractions (**D**), and tBuPhoX samples with or without snap-freezing of nuclei prior to crosslinking (**E**). Numbers indicate unique and shared residue pairs between datasets. **F**, Gene Ontology (GO) biological process enrichment analysis of proteins uniquely identified in the insoluble BSPNO2 fraction. Shown are the top enriched terms ranked by significance. **G**, Protein abundance distribution of crosslinked proteins in C. elegans nuclei. Proteins identified in the linear proteome search are shown in grey, whereas proteins supported by crosslinks are highlighted in blue. Protein abundance is ranked by LFQ intensity. **H**, Abundance-ranked protein distribution highlighting proteins involved in RNA-related pathways among crosslinked proteins. Grey, proteins identified in the linear proteome search; blue, proteins supported by interprotein crosslinks; red, selected proteins assigned to RNA-associated pathways. Highlighted proteins were selected based on abundance and annotation. **I**, Effect of precursor-intensity fractionation on crosslink identification. Mean URPs obtained from duplicate control injections without fractionation (blue), low-intensity precursor selection (orange), and high-intensity precursor selection (green). **J**, Overlap of URPs identified by the three acquisition strategies shown in **I**. Numbers indicate unique and shared URPs between acquisition methods. For the bar plots, the mean number of unique residue pairs is indicated within each bar. Individual technical replicates are shown as black dots. For SEC fractionation, individual fractions were combined to obtain one mean value.

For BSPNO2, the soluble fraction resulted in 1019 URPs, whereas the insoluble fraction yielded 373 URPs. Separate analysis of the insoluble fraction increased the total number of observed crosslinks by 18%, adding 222 newly identified URPs to a total of 1241 URPs (Figure 6C). In contrast, for SDA the soluble fraction yielded 62 URPs, whereas the insoluble fraction produced only 13 URPs. Although this added 12 newly identified crosslinks, the overall gain was only 16% (Figure 6D). Thus, the benefit of analyzing the insoluble fraction depends on crosslinker chemistry. BSPNO2 exhibits intermediate hydrophobicity (cLogP −1.0838) and high topological polar surface area (tPSA 199.60), whereas SDA is more hydrophobic (cLogP 0.5558) with lower polar surface area (tPSA 88.40)[75]. We hypothesize that the strong contribution of the soluble and insoluble fraction to the BSPNO2 dataset likely reflects preferential crosslinking of proteins within dense, poorly extractable nuclear assemblies such as chromatin- or scaffold-associated complexes. The balanced polarity of BSPNO2 may facilitate access to aqueous but insoluble protein environments and efficient lysine crosslinking, whereas the more hydrophobic photo-crosslinker SDA predominantly captured proteins already recovered in the soluble fraction. BSPNO2 displayed remarkable resistance to NHS-ester hydrolysis over a 24 h period (Supplementary Figure S13C). The MS1 signal of intact BSPNO2 decreased by only 1% after 1 h in anhydrous acetonitrile and by 3.7% after 24 h, whereas the corresponding hydrolysis product accumulated gradually over time. These results demonstrate that BSPNO2 remains largely intact under typical experimental handling conditions.

When using the tBuPhoX crosslinker, we performed, in addition to IMAC enrichment and SEC, experiments to evaluate the effect of snap-freezing prior to the crosslinking reaction. As expected, a single snap-freeze cycle of the nuclei increased the overall number of identified crosslinks, yielding 473 URPs, whereas the non-frozen sample resulted in 368 URPs. Notably, the insoluble fraction of the tBuPhoX no-snap-freeze condition contributed 101 URPs, which must be considered in the total number of crosslinks identified without prior freezing. Nevertheless, snap-freezing before crosslinking substantially increased the total number of detectable crosslinks. Both strategies also produced distinct subsets of unique crosslinks, with 137 URPs exclusive to the non-frozen condition and 242 URPs exclusive to the snap-frozen condition (Figure 6E). We therefore conclude that snap–freezing and no–snap–freezing can be used as complementary techniques to increase the overall number of crosslinks when studying nuclei interaction networks. The crosslinking experiment using LCSDA and DSSO in the nuclear environment yielded only 39 and 26 crosslinks in total, respectively. We were also interested in the proportion of protein–protein interactions (PPIs) identified in each experiment. For BSPNO2, we identified between 43 PPIs in the insoluble fraction and 183 PPIs in the soluble fraction, representing the best overall performance in terms of PPI detection among all crosslinkers tested. This was followed by tBuPhoX, with 118 PPIs in the worm entrapment dataset, 59 PPIs in the snap-frozen condition, 66 in the non-frozen condition, and 27 in the insoluble fraction. Notably, although not unexpected, the proportion of PPIs was higher in the non-snap-frozen sample than in the snap-frozen condition (Figure 6B). SDA experiments resulted in 26 PPIs in the soluble fraction and 11 in the insoluble fraction, whereas LCSDA and DSSO experiments yielded 29 and 26 PPIs, respectively. DSSO crosslinking showed in this experiment only PPI crosslinks.

We further examined differences in protein composition between the soluble and insoluble fractions by Gene Ontology (GO) enrichment analysis of biological processes (Figure 6F). Shown are the 15 most differentially enriched biological processes in the insoluble fraction of BSPNO2, with proteins involved in nucleobase, nucleic acid, RNA localisation and transport, nuclear export, mRNA transport, protein localisation as well as translational initiation and oogenesis. This is consistent with the hypothesis that BSPNO2 preferentially crosslinks nuclear assemblies such as chromatin–associated or RNA–bound complexes. Furthermore, we examined the abundance range of proteins for which crosslinks could be identified (Figure 6G). Crosslinks, including both inter– and intraprotein links, were detected across nearly the entire protein abundance range, extending to low LFQ intensities of approximately 10^5^. When considering only protein–protein interaction links, coverage became sparser, particularly at lower abundance levels, but still included proteins with intensities between 10^4^ and 10^5^ (Figure 6H). The practical detection limit for crosslinked peptides under the present workflow appears to be around LFQ intensities of 10^5^, consistent with the known abundance bias and reduced sampling efficiency of low-stoichiometry crosslinked species in complex mixtures[75, 76]. While reflecting on crosslinking abundances during mass spectrometry measurements, we designed a new strategy of crosslinked peptide enrichment during MS1 acquisition based on the observation that was described earlier[55], Supplemental Figure 8. This strategy is based on precursor-intensity selection for subsequent MS2 fragmentation. One injection was performed using an intensity threshold ranging from a minimum of 5,000 to a maximum of 10^5^, referred to as the low-intensity method (LowInt). A second injection used a filter ranging from 10^5^ to 1 × 10^20^, referred to as the high-intensity method (HighInt). These two injections were compared with a control method without intensity fractionation, using only a minimum threshold of 5,000 in duplicate injections to account for the equal number of technical analyses (Figure 6I). Duplicate control injections yielded, on average, 387 URPs, whereas the LowInt method identified 635 URPs and the HighInt method 732 URPs. Splitting precursor selection into two intensity ranges for MS2 fragmentation resulted in an overall gain of 33% in crosslink identifications, with 312 URPs unique to LowInt, 431 unique to HighInt, and 119 shared between both methods (Figure 6J). This gain could likely be further optimized by using narrower intensity-fractionation windows, although such approaches are limited by sample availability, as each injection required 250 ng of crosslinked material. This strategy holds strong potential to overcome current limitations of in vivo crosslinking workflows by improving crosslink identifications primarily at the mass spectrometry level rather than through further optimization of the crosslinking reaction itself.

Combining all crosslinker experiments resulted in an overall nuclear interaction map comprising 1822 intralinks and 463 protein–protein interactions across 755 proteins including ambiguous links. When ambiguous links are removed the numbers drop to 1013 intralinks, 241 PPI across 650 proteins. This interaction network clearly shows that each crosslinker contributed unique PPIs based on its distinct chemical properties, while a subset of PPIs was supported by multiple crosslinkers, increasing confidence in their detection and improving the robustness of the dataset. The interaction network is spanning protein complexes associated to for example U4/U6xU5 tri-snRNP complex, Box C/D RNP complex, Box H/ACA sno-RNP complex, Processosome, ribosomal complexes, messenger ribonucleoprotein complex, endoplasmatic complexes and the tRNA synthetase complex. Here, we do not focus on the biological interpretation of this dataset, but instead proceed with entrapment experiments using a subset of the complete interaction network (Supplemental Figure S8).

### 2.6 Entrapment benchmarking reveals how spectral-library design affects DIA-QCLMS confidence control

For entrapment experiments, we used experimentally generated *C. elegans* nuclei datasets crosslinked with BSPNO2 and tBuPhoX, together with a publicly available nuclei crosslinking dataset generated using DSG (PRIDE: PXD055488). The PhoX-crosslinked Cas9 sample was used to construct the target spectral library, including target–target (TT) entries as well as decoy–decoy (DD), target–decoy (TD), and decoy–target (DT) spectra. The respective entrapment libraries consisted of TT entries and the same decoy classes. Target and entrapment libraries were combined into a single composite library for DIA-based crosslink quantification. For Spectronaut analysis, default decoy generation was disabled to ensure use of the externally supplied library decoys. To assess target and decoy score distributions, quality-control filters were disabled and decoy identifications were exported through the report function. The exported results were then annotated with complimentary information from the spectral library such as the specific target-decoy labels for every peptide using a custom post-processing python script (as further described in section Rescoring of quantitative DIA crosslinking data).

False discovery rate (FDR) control in DIA-based crosslink quantification was systematically evaluated using different spectral-library configurations in combination with experimental entrapment strategies (Figure 7).

**Figure 7:**
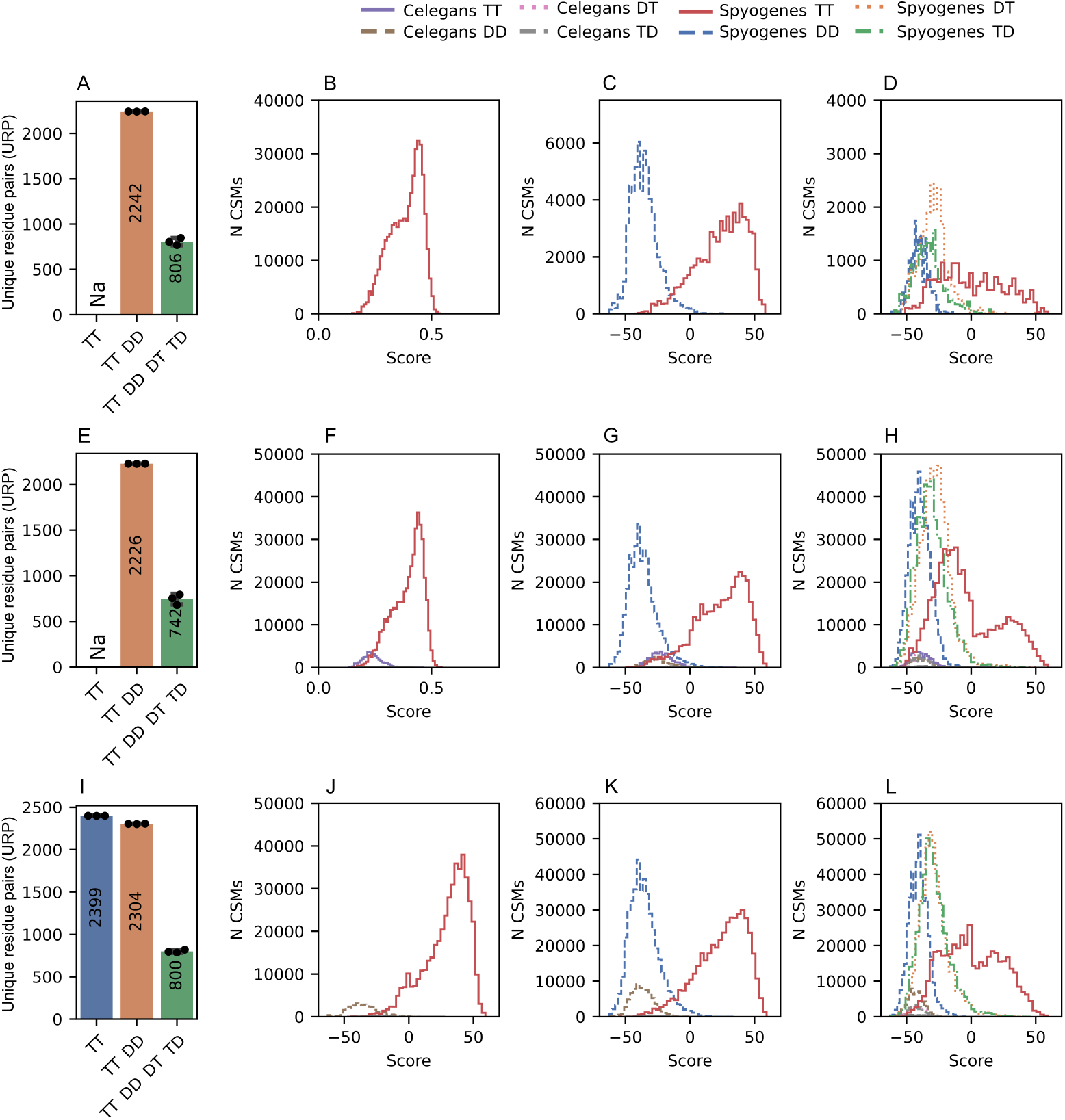
Spectral library FDR assessment using decoy spectra and entrapment strategies. Evaluation of spectral library false discovery rate (FDR) control using different decoy library configurations and entrapment strategies. **A**, **E**, **I**, Number of unique residue pairs (URPs) after quality assessment and filtering at a q-value of 0.01 (1% FDR). Crosslink identifications are shown for three spectral library configurations: target–target only (TT, blue), target–target with an additional decoy–decoy library (TT DD, orange), and a full target–decoy library containing TT, DD, DT, and TD classes (green). Results are shown for a library without entrapment spectra (**A**), with a C. elegans entrapment library using DSG crosslinking (**E**), and with a C. elegans entrapment library in which C. elegans entries were assigned as decoys (**I**). **B–D**, Crosslink spectrum match (CSM) score distributions for S. pyogenes library entries, showing target–target (TT, red), decoy–decoy (DD, blue), target–decoy (DT, green), and decoy–target (TD, yellow) identifications. Scores correspond to the Spectronaut Cscore. **F–H**, CSM score distributions including C. elegans entrapment hits. Target–target, decoy–decoy, target–decoy, and decoy–target classes are shown in violet, brown, grey, and pink, respectively. In this configuration, C. elegans entries were treated as target–target entries in the spectral library. **J–L**, Same analysis as in **F–H**, except that C. elegans entries in the entrapment library were defined as decoys. The entrapment library therefore contained only DD, DT, and TD classes for C. elegans sequences. To visualize decoy score distributions from Spectronaut outputs, all quality filters were removed and the “No Decoy” option in the report perspective was disabled. For bar plots, the mean number of unique residue pairs is indicated within each bar. Individual technical replicates are shown as black dots. Error bars (triplicates) represent the standard deviation calculated from replicate measurements. CSM score distributions were visualized as frequency histograms generated in Python using Matplotlib and Seaborn. Histograms were plotted with fixed-width bins (n = 40) and rendered as step outlines. No data transformation or statistical hypothesis testing was applied; plots represent descriptive comparisons of score distributions across groups.

Three library formats were compared: TT only, TT supplemented with DD entries (TT+DD), and a full target–decoy library containing TT, DD, DT, and TD classes (Figure 7A-D). In a second approach, the *C. elegans* entrapment library was included using the same three configurations (TT only, TT+DD, and full target–decoy; Figure 7E-H). Finally, a third strategy was tested in which all *C. elegans* TT entries were reclassified as decoys (Figure 7I-L) to evaluate the inflation of the decoy space to six fold. Using the standard *S. pyogenes* Cas9 benchmark library without entrapment entries and filtering at 1% q-value, the TT-only configuration yielded no quantified crosslinks, as FDR estimation cannot be performed without decoy entries. The TT+DD library resulted in 2,242 URPs, whereas inclusion of the full target–decoy model reduced identifications to 806 URPs (Figure 7A). This marked decrease suggests that mixed target–decoy precursor classes substantially increase score competition during library searching and impose more stringent confidence thresholds. Crosslink spectrum match score distributions showed clear separation between TT and DD matches, whereas DT and TD classes populated lower-scoring regions, consistent with expected false matches (Figure 7B-D).

We next introduced the *C. elegans* entrapment library as an orthogonal estimate of empirical error rates. When entrapment entries were treated as standard target sequences, identifications remained high for the TT+DD configuration (2,226 URPs) but decreased to 742 URPs for the full target–decoy model (Figure 7E). The TT+DD configuration produced five entrapment matches, corresponding to an empirical FDR of 0.2%. Score distributions showed that entrapment-derived matches partially overlapped with low-scoring target identifications, indicating that target-like entrapment entries can compete with true positives during scoring (Figure 7F-H). Nevertheless, true target distributions remained well separated from decoy-target and decoy classes. The reduction from 806 to 742 URPs is likely explained by expansion of the decoy search space from three classes (DD, DT, TD) to six classes (DD, DT, TD plus entrapment DD, DT, TD), suggesting slight overestimation of the nominal 1% FDR. Finally, all *C. elegans* entrapment entries were reclassified as decoys (TT → DD), restricting the entrapment space to DD, DT, and TD classes. Under this configuration, identifications were 2,239 URPs for TT only, with FDR estimation now enabled through the entrapment-derived DD entries, 2,242 URPs for TT+DD, and 800 URPs for the full target–decoy library (Figure 7I). Score distributions showed improved separation of true target matches from entrapment-derived decoy classes, enabling clearer interpretation of false-positive behavior (Figure 7J-L). These results indicate that assigning entrapment entries as decoys provides a cleaner framework for empirical FDR validation than treating absent-species sequences as targets. Across all tested approaches, full target–decoy libraries were consistently more conservative. This suggests that the threefold expansion of decoy space may slightly overestimate FDR, while simultaneously improving the specificity of crosslink spectrum matching. Overall, spectral-library composition strongly influenced DIA crosslink identifications and confidence control.

To assess the effect of entrapment library size, the *C. elegans* tBuPhoX entrapment dataset was reduced to 75%, 50%, and 25% of the original library size. This was achieved by downsizing the library while maintaining the distributions of peptide length, retention time, and precursor m/z comparable to those of the 100% library (Supplementary Figure S5). Crosslink spectral mapping in Spectronaut was performed as described above using only the full library configuration (TT, DD, DT, TD). Although target and decoy score distributions remained largely unchanged, and true target matches continued to separate well from decoy and entrapment classes, the number of quantifiable crosslinks at 1% FDR increased as library size decreased, from 498 URPs with the 100% entrapment library to 797 URPs with the 25% entrapment library (Supplemental Figure S5A-I). This gain is likely attributable to the reduced decoy search space, indicating that large entrapment libraries may lead to more conservative scoring and overestimation of FDR during spectral mapping. As the size of the decoy database increases, the probability that one or more decoy entries will randomly achieve high scores also increases[77] and leads in turn to less accepted CSMs.

To test different spectral library designs for entrapment experiments, we conducted two additional experiments: one using libraries with different crosslinker masses (altering only crosslinker-containing fragment ions) and another using PhoX-crosslinked human or *E. coli* ribosomes. Differences in crosslinker mass did not result in efficient separation of target and decoy distributions and consequently yielded only 11 URPs for the BSP dataset using DSG as entrapment, and 12 URPs for the inverse configuration (Supplementary Figure S6J-L). Using the same protein complex but from different organisms resulted in 8 URPs when *E. coli* ribosomes were used as the entrapment set, and 21 URPs when human ribosomes were used as entrapment. Although the entrapment score distributions showed partial separation from the true target CSMs, the DT, TD, and DD distributions of the target organism were poorly separated from true target matches. As a result, only a small number of target identifications passed the quality filters (Supplementary Figure S6J, M-P). Although the two previously mentioned entrapment strategies failed to sufficiently separate target and decoy distributions, incorporating all decoy types (DD, DT, TD) outperformed the TT-DD method for our initial entrapment experiment. This led to better distribution separation, more accurate FDR estimation for quantitative crosslinking data, and improved quality filtering in Spectronaut.

### 2.7 Empirical validation identifies Spectronaut Qvalue filtering as the most robust strategy in the tested workflow

Rescoring is the process of calculating a new match score based on additionally calculated or predicted features and has become the gold standard in linear peptide proteomics because it allows a more efficient separation of correct and incorrect peptide spectrum matches (PSMs)[78]. However, due to the more complex nature of both mass spectra and validation in crosslinking, rescoring is not yet as widely established. We have explored the process of rescoring for our quantitative DIA crosslinking workflow as well, using the output generated by Spectronaut as a basis (depicted in Figure 8). Spectronaut scores CSMs as one complete entity similar to how it would score a single peptide because there is no dedicated support for crosslinked peptides. Essentially we are using a workaround with our custom generated spectral library to enable crosslink identification, yet this employed scoring approach has the drawback that it does not account for the potential differences in match quality of the two individual crosslinked peptides. For example, it is not uncommon that one of the two peptides fragments well and forms many peaks in the resulting mass spectrum which in turn leads to a high identification score for that peptide. Contrary, the second peptide might not fragment at all but because its mass matches to the precursor in combination with the first peptide it is assigned to the CSM. Furthermore, because of the highly confident match of the first peptide, the CSM might also get a high score – this is however not desired because we do not know anything about the second peptide. In order to make this distinction we calculate several new features and scores based on the Spectronaut result file and the spectral library. Some of these features and scores are as follows: the number of matched ions per peptide, a relative score for each peptide by multiplying the CSM score with the fraction of matched ions to total ions per peptide, sequence coverage per peptide, the UniScore[79] for each peptide, the number of fragments containing crosslink modifications per peptide, and many more. An exhaustive list is given in the project’s GitHub repository (see section Code Availability). We then explored rescoring with two different approaches with a) either rescoring the complete CSM (from here on referred to as CSM-level rescoring), or b) splitting the CSM into two PSMs for each crosslinked peptide, then rescoring both PSMs separately and ultimately assigning the CSM score as the minimum of the two PSMs scores (from here on referred as PSM-level rescoring). Rescoring was performed using Mokapot[80] and emulating the Percolator model[81]. We then conducted the FDR estimation based on our newly calculated scores and tested if any of the new scores yielded an increase in the number of URPs at the same target FDR (i.e. 1% residue pair-level FDR). We also assessed the FDR estimation by analysing the number of false positives using entrapment data.

**Figure 8:**
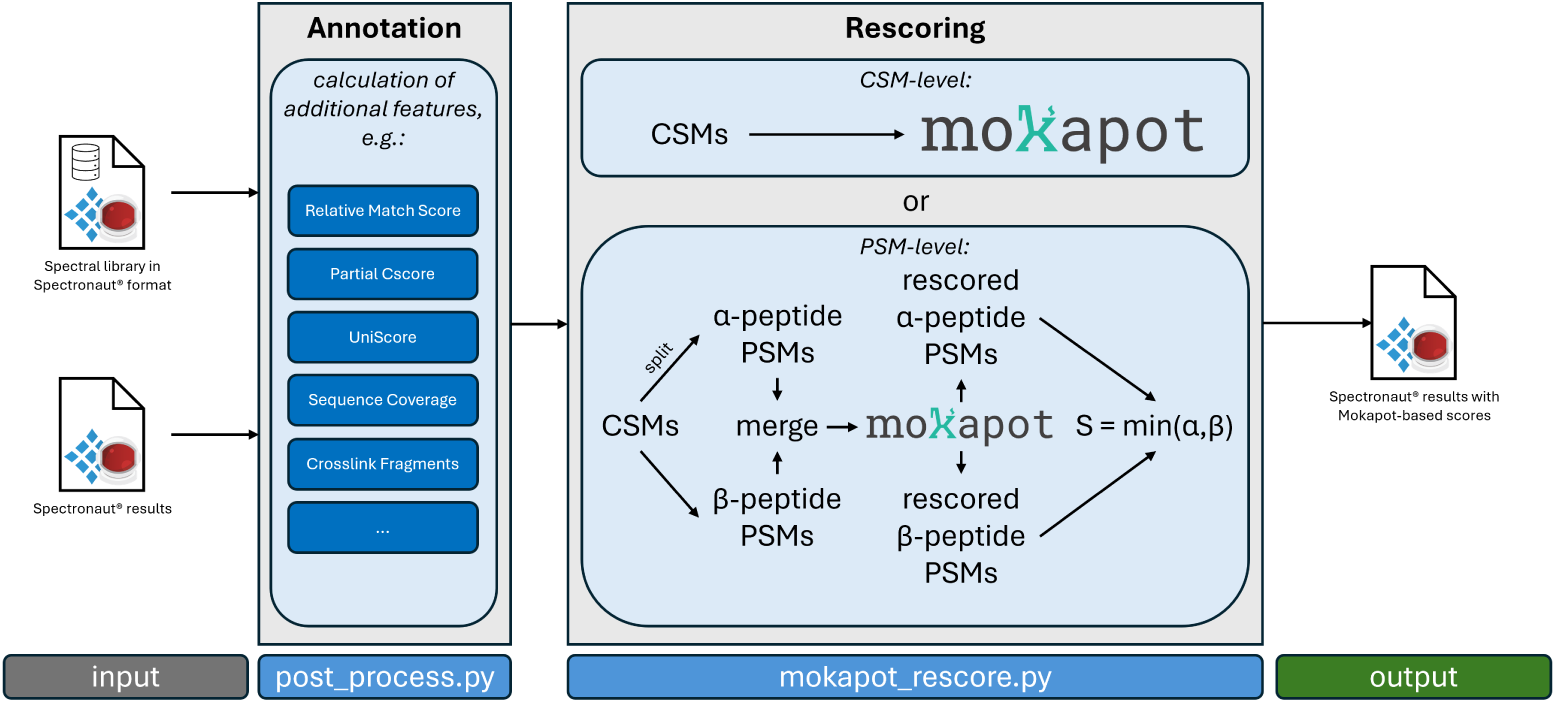
Schematic overview of the crosslink rescoring workflow. Our rescoring pipeline consisted of two steps: First the annotation of additional features based on the spectral library, for example the UniScore [79] and sequence coverage, and then rescoring with Mokapot [80]. For rescoring we pursued two different directions, either rescoring the crosslink-spectrum-matches (CSMs) directly using CSM specific features (CSM-level rescoring), or alternatively splitting the CSMs into peptide-spectrum-matches (PSMs) using the individual crosslinked alpha and beta peptides and rescoring based on PSM specific features (PSM-level rescoring). Rescored PSMs were then merged to CSMs again using the minimum of the two PSM scores as the CSM score. Both pipeline steps are implemented as standalone python scripts.

Generally we observed that CSM-level rescoring did not work, even though the new scores perfectly separated target and decoy hits leading to the highest number of URPs at 1% estimated FDR, this approach was severely underestimating the empirical (entrapment) FDR which was at 27% on average across the three replicates (Supplementary Figure S14A). This is also reflected in the score distributions shown in Supplementary Figure S14E-G demonstrating an accumulation of all decoy hits at the same score for our target system and entrapment system. The target-decoy score distribution of our target and entrapment system however does not efficiently separate entrapment from target system hits, leading to this high empirical FDR (Supplementary Figure S14G). Contrary, the FDR estimation with scores learned by PSM-level rescoring worked well at an empirical FDR of 0% - denoting that not a single entrapment hit was reported at 1% estimated FDR for all three replicates. This however came at the cost of more than 60% of the unique residue pairs being lost in comparison to the CSM-level rescoring approach (Supplementary Figure S14A). Nonetheless, score distributions also look visibly better with a more distinct separation of target and decoy hits as well as between our target and entrapment system (Supplementary Figure S14B-D). While this is a sizable improvement over the standard Spectronaut Cscore it does not outperform Spectronaut’s own validation (the Spectronaut Qvalue) which reports more than 2.5-times as many URPs at the same empirical FDR and also not reporting a single entrapment hit. In summary, this indicates that - despite not accounting for the intricate behaviour of crosslink validation - Spectronaut provides the best balance of target recovery and empirical entrapment control on its own without the need of any external validation or special extra steps - at least for our data. We therefore did not pursue this further for any of the other experiments in this manuscript.

### 2.8 State-resolved DIA-QCLMS captures ligand-dependent structural changes in UAP56

UAP56 (DDX39B) is a DExD-box ATP-dependent RNA clampase composed of two RecA-like domains (RecA1 and RecA2) connected by an interdomain linker[51, 52, 82]. Its catalytic core contains conserved sequence motifs that coordinate ATP binding, hydrolysis and RNA interaction. Motif I (Walker A) and motif II (DExD motif, Walker B) mediate ATP binding and hydrolysis, whereas motifs Ia, III, IV, V and VI contribute to RNA binding and coupling of ATPase activity to RNA engagement[82, 83]. Structural studies have shown that, in the absence of ligands, the two RecA domains adopt an open conformation, whereas ATP and RNA binding induce closure of the helicase core, bringing conserved motifs from both domains into proximity to form a composite active site[50, 82]. To define how human UAP56 engages RNA during its ATPase cycle, we established a state-resolved crosslinking mass spectrometry workflow using two complementary photoactivatable reagents, SDA and LCSDA, under conditions designed to capture either an apo/open conformer or an ATP/RNA-bound clamped conformer. Recombinant UAP56 was analysed either in the absence of ligands or in the presence of ATP/Mg^2+^ and polyuridine RNA (15U), conditions previously shown to promote closure of DExD-box helicases around RNA. Crosslinking reactions were benchmarked by SDS–PAGE and yielded robust state-dependent adduct formation with both chemistries (Supplementary Figure S3A,B). Due to the 89.4% sequence identity[84] and conserved helicase architecture between UAP56 and its paralogue DDX39A, many crosslinks were assigned ambiguously to both proteins. These shared-sequence matches were excluded from structural analyses as they do not represent true heterodimerization (Supplementary Figure S3C).

We next asked whether intramolecular restraints were consistent with the expected conformational transitions of UAP56. Mapping high-confidence residue-pair crosslinks onto AlphaFold3[85]-derived models of the open (Supplementary Figure S3H) and clamped (Supplementary Figure S3I) conformations revealed marked state specificity. SDA crosslinking identified 27 unique residue pairs (URPs) in the open state and 3 in the clamped state, whereas LCSDA identified 30 URPs in the open state and 25 in the clamped state. While SDA detected only a small subset of residue pairs enriched in the clamped state, LCSDA produced broadly similar crosslinking patterns under both conditions. This likely reflects the distinct structural sensitivity of the two reagents: SDA preferentially captures short-range (3.9 Å spacer) and orientation-dependent contacts formed upon domain closure, whereas the longer effective reach of LCSDA (12.5 Å spacer) accommodates residue pairs that remain proximal across conformational states. As expected, ATP/RNA binding appears to remodel local interdomain geometry without substantially altering the overall RecA-domain scaffold (Supplementary Figure S3D–G). The apo/open state of UAP56 likely represents a dynamic conformational ensemble rather than a single defined structure. Structural studies show that RecA1 (residues 45–257) and RecA2 (258–428) are connected by a flexible interdomain region spanning residues 251–264, enabling hinge-like rotation of RecA2 relative to RecA1[49, 82]. In the absence of ligands, this mobility can generate multiple transient domain orientations, explaining the dispersed crosslinking pattern observed in the open state. Residues within RecA1 motifs I (119–126) and II (221–224) can therefore come into transient proximity with RecA2 surfaces across different substates. This interpretation is supported by crosslinks exceeding theoretical distance constraints (Supplementary Figure S3J). In the open state, SDA- and LCSDA-derived restraints included overlength crosslinks with median *Cα* - *Cα* distances of 31.61 Å (SDA) and 25.89 Å (LCSDA), exceeding or approaching linker cutoffs. In contrast, clamped-state crosslinks remained largely within allowable ranges, with median distances of 16.26 Å (SDA) and 18.25 Å (LCSDA). Thus, ATP and RNA stabilize a clamped conformation in which both RecA domains close around RNA and align catalytic motifs into a defined active centre[50, 82]. Consistently, the clamped state yields fewer but more specific crosslinks, reflecting reduced interdomain flexibility. This is further supported by structural measurements showing that distances between Glu70–Leu262 (here Glu95–Leu287) and Pro173–Gln323 (here Pro198–Gln348) decrease from 27.8 Å and 24.4 Å in the open state to 18.9 Å and 16.1 Å in the clamped state[84].

Comparison of crosslinks between open and clamped states revealed only one clamped-state-specific crosslink in the SDA dataset and eleven in the LCSDA dataset (Figure 9B,C). Notably, clamped-state crosslinks were confined to sequence regions downstream of motif I (residues 119–126) and motif Ia (residues 147–152), whereas the N-terminal domain and the N-terminal portion of RecA1 remained largely uncrosslinked. This redistribution likely reflects ATP/RNA-induced domain closure, which buries or rigidifies N-terminal RecA1 surfaces while promoting defined contacts within the catalytic core. In addition, ligand-induced ordering of flexible regions, including the N-terminal extension, interdomain linker and RNA-contacting loops, may suppress heterogeneous crosslink formation and favour a more defined clamped architecture[50, 52, 82].

**Figure 9:**
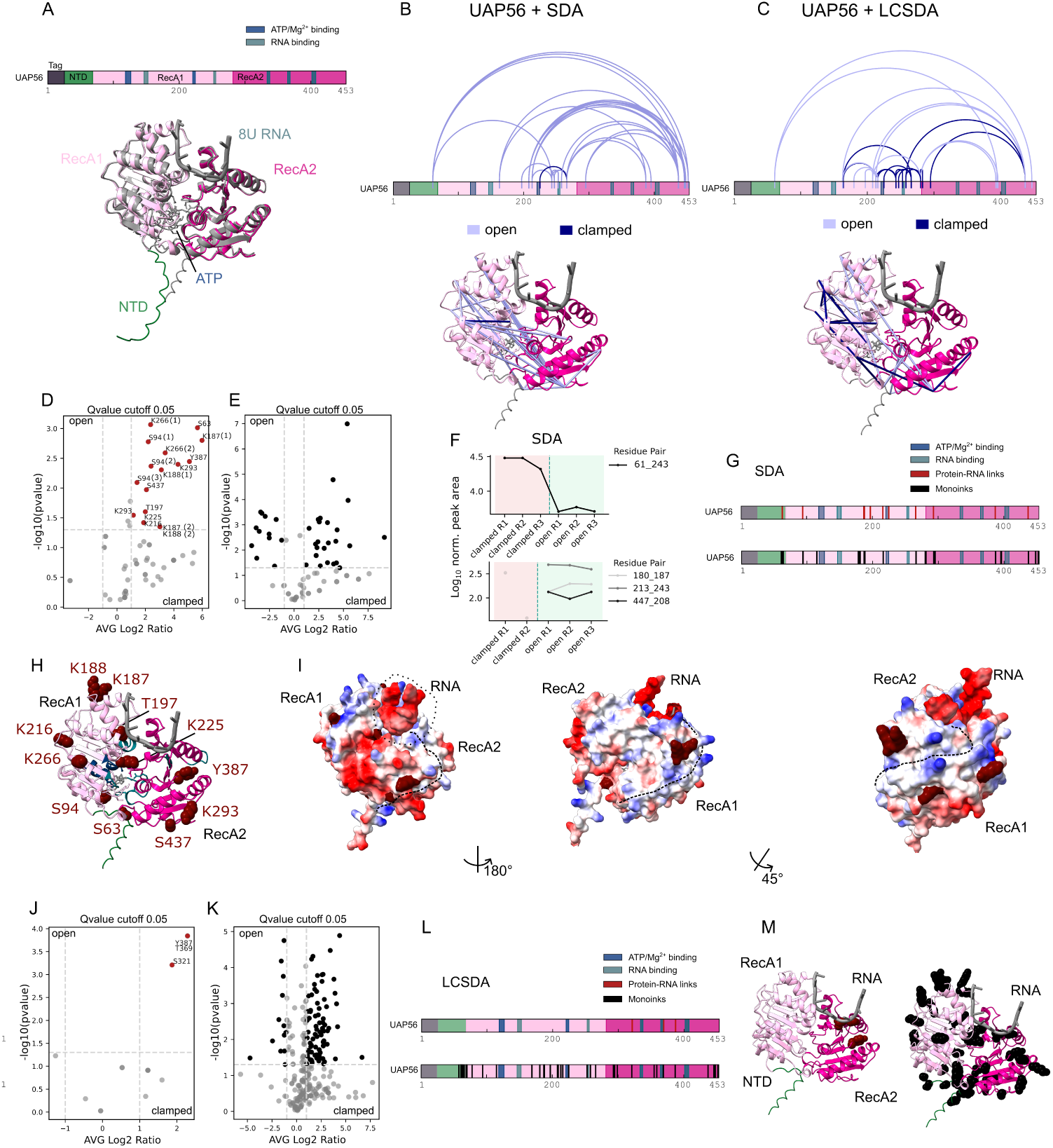
Structural dynamics and RNA binding of UAP56 using SDA and LCSDA crosslinking. Quantitative structural analysis of UAP56 in the absence (open state) or presence (clamped state) of ATP and 15U RNA using SDA and LCSDA crosslinking. **A**, Domain organization of recombinant UAP56 visualized in xiView[117]. The construct contains an N-terminal 10×His-3C tag (residues 1–25, black), followed by the N-terminal domain (26–68, green; endogenous residues 1–44), RecA1 lobe (69–281, light pink; endogenous residues 45–257), and RecA2 lobe (282–453, pink; endogenous residues 258–428). ATP/Mg^2+^-binding motifs (motif I, residues 119–126; motif II, residues 221–224) are shown in dark blue, whereas RNA-binding motifs (Ia, residues 147–152; III, residues 253–255; IV, residues 334–337; V, residues 365–368; VI, residues 402–407) are shown in pastel green. Residue numbering reflects the recombinant construct (25-residue N-terminal offset relative to endogenous UAP56). Below, a superposition of AlphaFold3-predicted open (coloured) and clamped (grey) conformations is shown; the clamped model includes ATP and 15U RNA (truncated to 8U RNA for clarity). **B**, Overlay of unique crosslinks identified with SDA in the open (light violet) and clamped (dark blue) states mapped onto the sequence and projected onto the AlphaFold3 clamped model. **C**, Equivalent analysis using LCSDA crosslinking. **D**, Volcano plot of quantitative protein–RNA crosslinks (SDA). Crosslinks significantly enriched in the clamped state are highlighted in red and labelled by residue. **E**, Volcano plot of quantified monolinks (SDA). Significantly enriched sites in the clamped state are shown in black. **F**, Quantitative summary of conformationally responsive crosslinks identified by SDA crosslinking. Differentially abundant links of the clamped conformer are shown at the top and links unique to the open conformation at the bottom. **G**, Sequence mapping of significant SDA-derived protein–RNA crosslinks (red) and monolinks (black) across structural domains. **H**, Clamped-state structural model with enriched protein–RNA crosslinked residues shown as red spheres. **I**, Electrostatic surface representation of the clamped model (red, negative; white, neutral; blue, positive). Enriched RNA-contact residues are shown in red. Three orientations illustrate the putative RNA-binding trajectory (RNA depicted as a dotted line). **J**, Volcano plot of quantitative protein–RNA crosslinks (LCSDA). **K**, Volcano plot of quantified monolinks (LCSDA). **L**, Sequence mapping of significant LCSDA-derived protein–RNA crosslinks (red) and monolinks (black). **M**, Structural mapping of LCSDA-enriched residues on the clamped model (protein–RNA crosslinks in red and monolinks in black). Volcano plots were generated using Seaborn scatterplots after quantitative analysis with the PTM Candidate workflow in Spectronaut (v. 20.4.260109.92449). Differential abundance between open and clamped states was calculated using Spectronaut’s default two-sided t-test with Benjamini–Hochberg multiple testing correction. Volcano plots display log_2_ fold change versus − log_10_(q-value), and significance was defined as p < 0.05 with an additional quality filter of Q-value < 0.05 (**D**, **E**, **J**, and **K**).

We next used UV-induced protein–RNA crosslinking to identify residues that directly contact RNA. DIA data were analysed in Spectronaut using the directDIA workflow and filtered at 1% q-value. Site localization was performed using the Spectronaut PTM localization algorithm. Quantitative analysis of SDA-derived RNA adducts identified a discrete set of residues significantly enriched in the clamped state (Figure 9D), clustering across both RecA domains and the interdomain interface. Residues K266, K188, K187 and S94 were supported by multiple precursor species, increasing confidence in site assignment. In total, 33 monolink sites were enriched in the clamped state and 14 in the open state. Mapping of RNA adducts revealed clustering around motif II (221–224), motif Ia (147–152) and motif VI (402–407). Additional protein–RNA links were detected within the Q motif region (65–72; here 90–97), which is associated with ATP binding[84]. We note that certain sites (e.g. S63/S94) may reflect nucleotide-derived adducts rather than direct RNA contacts and are therefore interpreted as nucleotide-proximal. A prominent RNA-contact cluster was observed at K187/K188 and T197, suggesting a RecA1-proximal binding region. Additional sites at K216 and K225 localize adjacent to motif II, placing RNA contacts in close proximity to the catalytic ATPase center. Sites K266 and K293 likely report RNA positioning from RecA1 across the interdomain cleft towards RecA2, whereas Y387 and S437 extend along the RecA2 surface. Mapping onto electrostatic surfaces revealed alignment of RNA-contact residues with positively charged patches, consistent with RNA backbone recognition and suggesting a continuous RNA-binding trajectory (Figure 9I). Notably, T197 and K225 correspond to RNA-contact sites observed in cryo-EM structure 7ZNK[62]. LCSDA yielded fewer RNA-crosslinked residues, identifying three enriched sites (S321, T369 and Y387) localized to the RecA2 region (Figure 9J–M). These overlap with the broader RNA-contact surface detected by SDA but likely reflect stronger or more persistent contacts due to differences in crosslinker geometry.

Monolink analysis was used as a proxy for conformation-dependent residue accessibility[86, 87] and revealed clear differences between the open and ATP/RNA-bound clamped states (Figure 9E,K). The open state showed enrichment at residues K225, K187/K188, K180, S155, T283, Y387 and S393, particularly near the interdomain cleft, indicating increased accessibility prior to closure. In contrast, clamped-state monolinks were more broadly distributed across peripheral regions, consistent with closure of the catalytic core and exposure of distal surfaces (Figure 9M). Quantitative analysis of crosslinked peptides identified only a limited number of state-responsive intramolecular restraints. For SDA, five crosslinks were quantified in the open state and seven in the clamped state; for LCSDA, seven were quantified in each condition. Among these, only a small subset showed significant changes. In the SDA dataset, crosslink 61 to 243 was enriched in the clamped state, whereas 180 to 187, 213 to 243 and 447 to 208 were enriched in the open state (Figure 9F). In the LCSDA dataset, crosslinks 188 to 187, 213 to 243 and 188 to 181 were enriched in the clamped state, whereas 155 to 181 was enriched in the open state (Supplementary Figure S7A,B).

Given the limited number of quantifiable crosslinks, likely reflecting the low abundance of crosslinked species and conservative FDR filtering, individual state-specific restraints should be interpreted cautiously. Although the identities of significant crosslinks differed between SDA and LCSDA, both datasets converged on the same overall structural model. Crosslinks detected in the apo state were more broadly distributed and frequently exceeded theoretical distance constraints, whereas ATP/RNA-bound samples showed fewer, more localized restraints that were largely compatible with the clamped conformation. These observations are consistent with structural studies of UAP56 and other DExD-box ATPases, which undergo ATP- and RNA-dependent closure of the two RecA domains into a compact catalytic state. We therefore interpret the observed crosslink changes as evidence for a transition from a conformationally heterogeneous open ensemble towards a structurally constrained clamped conformation upon ATP/RNA binding. Individual crosslinks highlight regions particularly responsive to this transition but should not be overinterpreted in isolation.

## 3 Discussion

### 3.1 Quantitative and empirically validated crosslinking mass spectrometry

Quantitative crosslinking mass spectrometry provides a powerful means to probe protein structure and dynamics in solution, yet its broader application has been limited by sparse sampling, missing values and challenges in confidence estimation for crosslinked peptide identifications. Here, we establish a data-independent acquisition (DIA)-based quantitative crosslinking mass spectrometry framework that addresses these limitations through the integration of optimized acquisition strategies, crosslink-aware spectral libraries and empirical false discovery rate validation. A central component of this work is the implementation of a crosslink-aware spectral library design that explicitly accounts for the paired nature of crosslinked peptides. By incorporating target–target, target–decoy, decoy–target and decoy–decoy entries, the workflow preserves the full target–decoy structure inherent to crosslinking data and enables empirical assessment of identification confidence in DIA analysis[53]. In combination with entrapment-based benchmarking, this strategy provides a practical framework for evaluating FDR behavior in complex datasets. Within the tested conditions, native Spectronaut Qvalue filtering yielded the most favorable balance between target recovery and empirical control of false positives, indicating that robust validation can be achieved without additional post hoc rescoring when appropriate spectral libraries and acquisition strategies are employed.

### 3.2 Optimization of DIA acquisition and quantitative performance

Our systematic evaluation of DIA acquisition parameters further revealed fundamental trade-offs that govern quantitative crosslinking experiments. Narrow isolation windows maximized crosslink identifications but reduced chromatographic sampling, whereas broader windows improved sampling density at the expense of precursor selectivity. Similarly, FAIMS compensation voltage stepping substantially increased precursor coverage but reduced quantitative precision owing to prolonged duty cycles and reduced data points per chromatographic peak. These observations demonstrate that DIA-QCLMS workflows must be optimized to balance identification depth and quantitative robustness rather than maximizing either parameter in isolation. The acquisition strategy established here therefore provides practical design principles for DIA-based quantitative crosslinking experiments and defines an experimentally validated compromise between precursor coverage, quantitative reproducibility and chromatographic sampling density. Crosslinked *Caenorhabditis elegans* nuclei were used both to generate crosslinker-specific spectral libraries and to establish entrapment datasets for empirical validation of identification confidence. The resulting DDA-derived datasets provide a realistic representation of crosslinking complexity in cellular systems and enable systematic evaluation of false-positive rates under heterogeneous biological conditions. In contrast, DIA measurements were specifically applied within an entrapment framework to assess identification performance and FDR behavior. These analyses demonstrate that accurate confidence estimation in DIA-QCLMS critically depends on spectral library composition and validation strategy, particularly in highly complex backgrounds.

### 3.3 Rescoring strategies and future integration of predictive spectral libraries

Crosslinked *Caenorhabditis elegans* nuclei were used to generate crosslinker-specific spectral libraries and to establish entrapment datasets for empirical validation of identification confidence. The resulting DDA-derived datasets provide a realistic representation of crosslinking complexity in cellular systems and enable systematic evaluation of false-positive rates under heterogeneous conditions. In contrast, DIA measurements were applied specifically within an entrapment framework to assess identification performance and FDR behavior. These analyses demonstrate that accurate confidence estimation in DIA-QCLMS critically depends on spectral library composition and validation strategy, particularly in complex backgrounds. Despite systematic evaluation of alternative validation strategies, we did not observe an improvement in identification performance using our rescoring pipeline. We attribute this to the strong baseline performance of Spectronaut’s native validation, which appears to effectively capture both linear and crosslinked peptide features within the tested datasets. This suggests that, under these conditions, additional rescoring based on features derived from spectral libraries and crosslink-spectrum matches (CSMs) provides limited incremental benefit. However, rescoring approaches may benefit from the inclusion of orthogonal, prediction-based features, such as retention time and fragment intensity predictions generated by tools including xiRT[88], Prosit-XL[89], or XL-MSDigger[90], which could provide complementary information not already captured by the native scoring framework. Notably, we were unable to achieve fully reproducible rescoring results using Mokapot[80], observing variations of approximately ±10% in identification counts across repeated runs with identical inputs and random seeds. While these fluctuations did not affect the overall conclusions of this study, as differences between validation strategies exceeded this range, they highlight potential stability limitations when applying machine learning-based rescoring approaches to crosslinking datasets.

In this study, quantitative analysis was performed using experimentally derived spectral libraries generated from prior DDA searches. This approach ensures accurate spectral representation of crosslinked peptides and enables robust DIA-based quantification. However, it also restricts analysis to crosslinks previously identified in DDA data, potentially excluding low-abundance or condition-specific crosslinks that may be detectable in DIA measurements alone. Future developments may address this limitation through hybrid strategies combining experimental and predicted spectral libraries, for example using Prosit-XL[89], or XL-MSDigger[90], thereby integrating the quantitative accuracy of experimental libraries with the broader coverage of prediction-based approaches. The exclusive use of predicted spectral libraries remains challenging in crosslinking applications. Many prediction frameworks require calibration using experimentally observed spectra, typically derived from DDA data, and therefore do not fully eliminate the need for prior identification workflows. In addition, the combinatorial expansion of crosslinked peptide pairs imposes practical limitations on search space generation, particularly for large proteomes, where exhaustive enumeration of all possible peptide pairs becomes computationally intractable. These considerations underscore the continued importance of experimentally derived libraries for quantitative crosslinking workflows, while highlighting opportunities for future integration of predictive approaches.

### 3.4 A multi-crosslinker *C. elegans* nuclear interaction resource

Beyond its role in method development, the crosslinked *Caenorhabditis elegans* nuclei dataset constitutes a multi-crosslinker protein–protein interaction resource that, to our knowledge, has not previously been reported for this system. By combining five complementary crosslinking chemistries, including SDA, LCSDA, DSSO, tBuPhoX and BSPNO2, the dataset captures a broad spectrum of interaction distances and chemical reactivities, enabling expanded structural coverage of nuclear protein assemblies. Notably, BSPNO2-based crosslinking yielded a distinct interaction profile enriched in RNA/DNA-associated complexes, suggesting that crosslinker chemistry can bias detection towards specific functional classes of proteins and highlighting the potential of tailored crosslinking strategies for probing defined sub-networks. This observation suggests that crosslinker chemistry strongly influences the detectable interaction landscape and indicates that BSPNO2 may be particularly well suited for probing ribonucleoprotein and chromatin-associated assemblies in situ. Recovery of crosslinks from detergent-solubilized insoluble nuclear fractions substantially expanded interaction coverage, particularly for RNA-associated complexes, indicating that structurally constrained or partially insoluble nuclear assemblies remain underrepresented in conventional soluble crosslinking workflows. Notably, the workflow enabled confident identification of inter-protein crosslinks from proteins with LFQ intensities below 10^5^, demonstrating sufficient sensitivity to interrogate low-abundance interaction networks in complex proteomes.

Furthermore, implementation of an MS1-intensity fractionation strategy increased crosslink identifications by approximately 33% per duplicate injection, demonstrating that substantial improvements in crosslink coverage can be achieved at the acquisition level without additional biochemical enrichment. This finding is particularly notable because it reflects improved utilization of instrument duty cycle and precursor sampling rather than optimization of sample preparation workflows. Collectively, these results demonstrate how the combination of orthogonal crosslinking chemistries and acquisition strategies can substantially improve both the depth and diversity of crosslinking datasets in complex biological systems. Consequently, the DDA-derived *C. elegans* dataset provides a valuable resource for benchmarking, method development and future biological investigations. In particular, the availability of a multi-chemistry crosslinking dataset within a single biological system creates opportunities to systematically evaluate crosslinker-specific biases, improve computational scoring strategies and potentially train predictive models for crosslink identification.

### 3.5 State-resolved structural analysis of UAP56 by DIA-QCLMS

As a proof-of-principle application, we applied the DIA-QCLMS workflow to the ATP-dependent RNA helicase UAP56 (DDX39B), a model system for ligand-dependent conformational rearrangements. The analysis captured changes in intramolecular crosslinks, monolink abundance and protein–RNA adducts between apo and ATP/RNA-bound states, consistent with the transition from an open to a clamped conformation. Rather than defining RNA-binding sites at residue-level resolution, these data identify candidate RNA-proximal regions and conformationally responsive elements within the helicase core. These findings illustrate how DIA-QCLMS can be used to compare structural states and generate hypotheses on protein–RNA interactions in systems characterized by conformational heterogeneity.

More broadly, integration of monolinks and protein–RNA crosslinks extends the analytical scope of quantitative crosslinking beyond pairwise residue restraints alone. Monolinks provide complementary information on local residue accessibility and conformational shielding, whereas protein–RNA adducts report on RNA-proximal environments and ligand-dependent interaction surfaces. The identification of state-specific monolinks and RNA adducts in UAP56 demonstrates how these orthogonal features can contribute to interpretation of conformational transitions and accessibility remodeling. Although interpretation of such features requires careful consideration of crosslinker chemistry, ligand proximity effects and detection biases, their combined analysis enables a more comprehensive view of structural remodeling across biological states.

### 3.6 Limitations and future perspectives

Several limitations of the current workflow should nevertheless be considered. First, crosslinked peptides remain low-abundance species, resulting in limited numbers of quantifiable features in some datasets and constraining statistical power. Second, assignment of protein–RNA adducts can be influenced by nucleotide-derived modifications or proximity to nucleotide-binding sites, particularly in the absence of ATP-only or RNA-only control conditions. Third, spectral library-based DIA workflows inherently bias quantification towards features identified in prior DDA experiments, potentially limiting discovery of novel crosslinks in DIA-only analyses. Finally, acquisition parameters and performance characteristics remain instrument- and sample-dependent and may therefore require optimization for different biological systems. Future developments will focus on improving sensitivity and coverage of crosslinked peptides through advances in enrichment methods, refinement of MS1-intensity fractionation approaches and integration of machine learning-based scoring strategies. In addition, combination with complementary structural proteomics techniques, including hydrogen–deuterium exchange and limited proteolysis, as well as development of time-resolved crosslinking strategies, may further enhance the ability to resolve dynamic conformational changes. Continued efforts towards standardized data formats, validation strategies and open-source analysis tools will be essential to facilitate broader adoption of DIA-QCLMS across the structural proteomics community. Collectively, these advances position DIA-QCLMS as a scalable and quantitatively robust platform for structural proteomics, capable of resolving interaction landscapes, conformational ensembles and ligand-dependent structural remodeling across complex biological systems.

## 4 Methods

### 4.1 Reagents

### 4.2 Crosslinking reaction for Cas9

Cas9–Halo protein was crosslinked using PhoX. The crosslinker was prepared as a 50 mM stock solution in dry acetonitrile (ACN). For crosslinking, Cas9 was diluted in 50 mM HEPES to a final protein concentration of 1 *µ*g *µ*l*^−^*^1^, and PhoX was added to a final concentration of 1 mM. Reactions were incubated on ice for 45 min and quenched by addition of Tris buffer to a final concentration of 100 mM.

### 4.3 In-Solution Digest of Cas9 QC samples

For in-solution digestion, Cas9 was reduced with 10 mM dithiothreitol (DTT) for 30 min at 50 *^◦^*C, followed by 10 min of water bath sonication, and subsequently alkylated with 50 mM iodoacetamide (IAA) for 30 min in the dark. Proteins were digested with trypsin at an enzyme-to-protein ratio of 1:100 (w/w) and incubated overnight at 37 *^◦^*C. Digestion was terminated by addition of trifluoroacetic acid (TFA) to a final concentration of 0.2

### 4.4 Chemical synthesis of the BSPNO2 (CLIP) crosslinker

The CLIP crosslinker was prepared according to Gao et al.[57–59] in three steps (see Figure M1). First, trinitrile a was hydrolyzed with HCl to give triacid b. Next, one of the three carboxyl groups was activated with EDC in a statistical manner and coupled with propargylamine to yield c. Finally, the two remaining carboxyl groups were activated with EDC and converted to NHS esters to afford the CLIP crosslinker.

**Figure M1:**
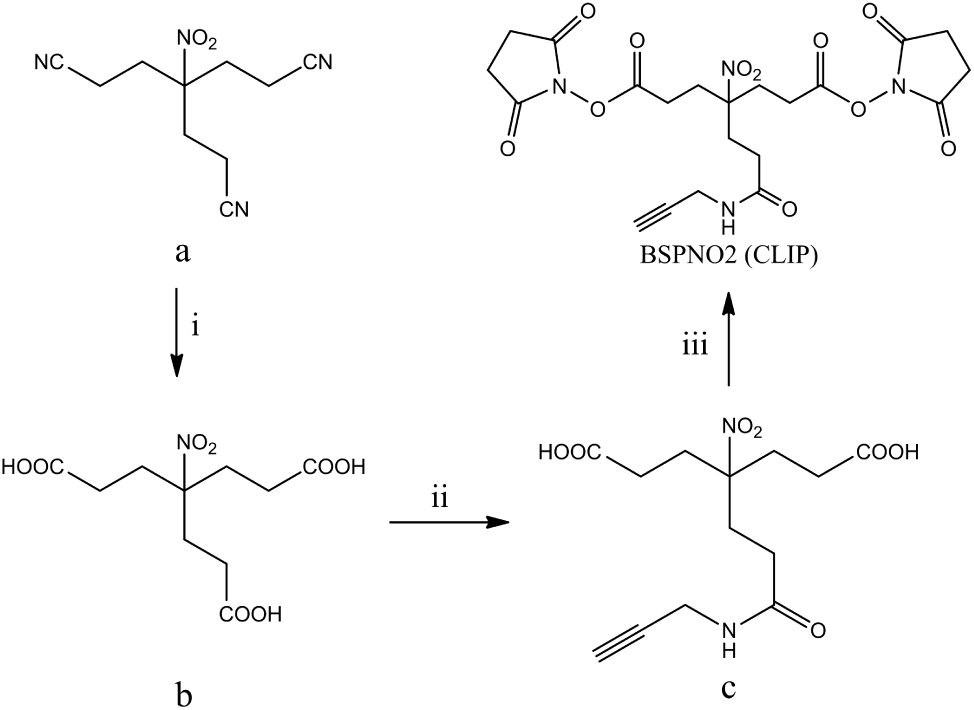
Synthesis of the BSP crosslinker. i: HCl, reflux; ii: EDC, propargylamine; iii: EDC, NHS.

### 4.5 Purification of the BSPNO2 (CLIP) crosslinker using HPLC

The crude CLIP crosslinker was dissolved in dry DMSO and cooled to −20 *^◦^*C prior to HPLC purification. The solution was maintained at low temperature throughout all purification steps. Crosslinker purification was performed on a Kinetex column (2.6 *µ*m, C18, 100 Å, 150 × 2.1 mm, part no. 00F-4462-AN, Phenomenex) using a 50 min gradient from 20 to 60% B (A: 0.1% TFA; B: ACN containing 0.1% TFA) at a flow rate of 0.1 mL min*^−^*^1^. Fractions were collected from 10 to 40 min and transferred to precooled low-bind reaction tubes. The ACN content was adjusted to a final concentration of 10%. Samples were snap-frozen in liquid nitrogen and lyophilised overnight. The final white fluffy powder of the BSPNO2 product could be identified in fractions eluting at and around 26 min. Representative MS_1_ spectra of the purified BSPNO2 crosslinker are shown in Supplementary Figure S13A,B.

### 4.6 Characterisation of the BSPNO2 crosslinker using mass spectrometry

To monitor hydrolysis over time, the BSPNO2 crosslinker was analysed repeatedly over a total period of 24 h. During the initial 9 h of the experiment, injections were acquired at approximately 9-min intervals. Beyond 9 h, the interval between consecutive injections was increased as required, resulting in variable sampling intervals. The fraction containing the BSPNO2 crosslinker (26 min) was dissolved in acetonitrile (ACN) prior to analysis and diluted 1:10 (v/v) with ACN. To minimise hydrolysis before measurement, the first injection was performed within 2 min of sample dissolution. A volume of 1 *µ*L was injected into an Ultimate 3000 HPLC system (Thermo Fisher Scientific, Germany) configured for flow-injection analysis and coupled directly to an Exploris 480 (Thermo Fisher Scientific) mass spectrometer (MS). No chromatographic separation was performed. An isocratic mobile phase consisting of 95% ACN and 5% 0.1% formic acid at a flow rate of 100 mL min*^−^*^1^ was applied. The MS was operated in alternating polarity mode. Electrospray ionisation (ESI) parameters were set as follows: spray voltage, +3.4/−2.8 kV; sheath gas, 35; auxiliary gas, 5; ion transfer tube temperature, 300 *^◦^*C; vaporiser temperature, 250 *^◦^*C. Data-dependent spectra were acquired over an *m/z* range of 290–700 at a resolving power of 120,000 (at *m/z* 200). The data were processed using Skyline (v25.1.0.237)[69, 70] and Microsoft Excel.

### 4.7 10×His-3C-UAP56 fusion protein purification

10×His-3C-UAP56 was purified as described previously[49, 50]. In short, 10×His-3C-UAP56 was expressed in Escherichia coli BL21(DE3) RIL cells grown in LB medium. Protein expression was induced at OD600 = 1.0 with 0.5 mM IPTG and cells were incubated at 37 *^◦^*C for 3 h. Cells were lysed by sonication in lysis buffer (25 mM HEPES pH 7.9, 5% glycerol, 300 mM NaCl, 20 mM imidazole, 0.05% Tween-20, protease inhibitors), and the lysate was clarified by centrifugation and filtration. The supernatant was purified by Ni^2+^ affinity chromatography (HisTrap HP, Cytiva), washed with 44 mM imidazole and eluted using a 50–300 mM imidazole gradient. Fractions were diluted to 100 mM NaCl and further purified by anion-exchange chromatography (HiTrap Q, Cytiva) with elution at 200–400 mM NaCl. Final purification was performed by size-exclusion chromatography (Superdex 200, Cytiva) in buffer containing 25 mM HEPES pH 7.9, 5% glycerol, 250 mM NaCl and 1 mM DTT. Purified protein was concentrated, flash-frozen and stored at −80 *^◦^*C.

### 4.8 Size-exclusion chromatography of UAP56–RNA–ATP complexes

Size-exclusion chromatography was performed using a Superdex 200 Increase 10/300 column (Cytiva) equilibrated in gel filtration buffer (25 mM HEPES pH 7.9, 100 mM NaCl, 1 mM MgCl_2_, and 1 mM TCEP). For complex formation, 10×His-3C-UAP56 (600 *µ*g) was incubated with 15U RNA (120 *µ*M) and ATP (1 mM) in a total volume of 600 *µ*L, with the final NaCl concentration adjusted to 100 mM. Prior to incubation, protein stocks were clarified by centrifugation (10 min, 4 *^◦^*C, maximum speed) to remove aggregates. The reaction was incubated at 24 *^◦^*C for 1 h. A total of 500 *µ*L of the sample was loaded onto the column using a 500 *µ*L loop (loop flushed with 2 mL equilibration buffer). The column was operated at a flow rate of 0.5 mL min*^−^*^1^, with 0.05 column volumes (CV) equilibration prior to injection and elution performed over 1 CV. Fractions were collected in 100 *µ*L increments starting after 0.2 CV.

### 4.9 Crosslinking of open and clamped UAP56

UAP56 (20 µM) in crosslinking buffer (25 mM HEPES pH 7.9, 100 mM KCl, 1 mM MgCl_2_) was incubated either with 1 mM ATP and 120 µM 15U RNA (clamped conformation) for 1 h at room temperature or without ATP and RNA (open conformation) prior to crosslinking. Crosslinking was performed using 1 mM SDA or 1 mM LCSDA for 30 min on ice to allow the NHS ester reaction. The reaction was quenched by addition of 20 mM Tris, followed by 25 s ultraviolet (UV) irradiation (365 nm, 100% intensity) using a HONLE LED CUBE 100 to induce photo-crosslinking. After UV irradiation, samples were incubated in Laemmli sample buffer containing 100 mM DTT for 10 min at 80 *^◦^*C and subsequently separated by SDS–PAGE to isolate monomeric species.

### 4.10 In-gel digestion of crosslinked 10×His-3C-UAP56 in open and clamped state

Samples were separated on Novex NuPAGE 1-mm 3-8% Tris–acetate gels (Invitrogen) using Tris–acetate running buffer (Invitrogen). Gels were fixed in 50% methanol and 5% acetic acid and stained using a colloidal Coomassie Blue kit (MBS Facility). Protein bands of interest were excised and transferred into low-binding tubes (Microtubes “Maxymum Recovery” Axygen). Gel pieces were destained with 30% (v/v) acetonitrile (ACN) in HEPES buffer pH 7.3 for 45 min; this step was repeated until the gel pieces were clear. Proteins were reduced with 100 mM dithiothreitol and alkylated with 55 mM iodoacetamide. Following reduction and alkylation, the supernatant was removed and gel pieces were dehydrated with 100% ACN. Proteins were digested with trypsin (300 ng *µl^−^*^1^). Peptides were extracted using 80% (v/v) ACN in 0.1% (v/v) TFA, desalted using C18 StageTips[92], and eluted with 80% (v/v) ACN, 0.1% (v/v) TFA prior to mass spectrometric analysis. Eluted peptides were concentrated in a vacuum concentrator (Eppendorf) and resuspended in 0.1% (v/v) Trifluoriformic acid (TFA), followed by 3 min sonication.

### 4.11 Preparation of the *C. elegans* nuclear extract and crosslinking

15 mL of *Caenorhabditis elegans* N2 Bristol worms were collected based on a published protocol[93]. Briefly, the worms were grown until the adult stage on NGM++ plates seeded with OP50. They were bleached, and the eggs were transferred to plates devoid of bacteria for hatching. The arrested L1 larvae were then transferred to a flask containing OP50 bacteria suspended in S-medium and were left in a shaking incubator for 3 days (20°C, 70rpm). Adult worms were then collected by filtering the worm culture through a 40µm mesh. Worms were washed off the mesh with ice-cold M9 supplemented with 0.01% Tween-20 followed by an additional wash. 1 ml of worms were resuspended with 3mL buffer containing 10mM HEPES-KOH (pH=7.6), 1mM EDTA (pH=8), 10mM KCl, 1.5mM MgCl2, 0.25mM sucrose, 1mM PMSF, and 1mM DTT. Then, samples were snap frozen in liquid nitrogen and stored at −80°C until further use. 4 ml of worms were thawed, dounced, and passed through filters of pore sizes 100µm and 40µm following this protocol[94]. The filtered solution was centrifuged at 300g for 2 minutes at 4°C. The supernatant was centrifuged at 2500g for 10 minutes at 4°C. The pellet thus obtained was resuspended using a buffer containing 20mM of HEPES, 40mM NaCl, 90mM KCl, 1mM EDTA, 1%(v/v) NP40 and 1mM PMSF and incubated at 4°C for 15 minutes. This solution was centrifuged and the pellet was resuspended in 50mM HEPES (pH 7.5), 50mM potassium acetate and 10mM magnesium acetate tetrahydrate for crosslinking. Germline nuclei were crosslinked using the following crosslinkers: SDA (60mM stock, 2x 2 mM final concentration), LC-SDA (60mM stock, 2x 2mM final cenc.), DSSO (80mM stock, 5mM final conc.), tBUPhoX (30mM, 2x 1mM final conc.) and BSPNO2 (80mM stock, 5mM final conc.) (tBuPhoX, Cat-no.: A52287, SDA, Cat-no.: 26167, LC-SDA Cat-no.: 26168, DSSO Cat-no.: A33545, Thermo Fisher Scientific, BSP[57, 59]). The stock solutions were prepared using dry ACN and the crosslink reaction was carried out for 1h on ice, following quenching by adding 100 mM Tris (pH 7.4) and incubating it for 10 minutes at room temperature.

### 4.12 In-solution digest of crosslinked nuclei

Isolated and crosslinked nuclei were solubilized in 8 M urea and 0.05% final conc. n-Dodecyl *β*-D-Maltosid (DDM), following cell lysis by sonication (UP100H Ultrasonic Processor, Hielscher) (30 s, 80% amplitude, 0.5 s cycle) with intermittent cooling on ice; this step was repeated three times. Proteins were reduced with 10 mM dithiothreitol (DTT) for 30 min at 50 °C and alkylated with 50 mM iodoacetamide (IAA) for 30 min at room temperature in the dark. The sample was diluted to 2 M urea using 50 mM HEPES (pH 7.3). To digest nucleic acids, MgCl_2_ (1 mM) and Benzonase (25 U *µl^−^*^1^) were added and the mixture was incubated for 1 h at 37 *^◦^*C. Proteins were subsequently digested with LysC (1:100, 1.5 h, 37 *^◦^*C), followed by trypsin digestion at a 1:50 (enzyme-to-protein, w/w) ratio overnight at 37 *^◦^*C. Digestion was terminated by acidification with trifluoroacetic acid (TFA) to a final concentration of 0.5%. Samples were centrifuged (20,000g, 10 min) to remove precipitates. The supernatant was thereafter labelled as soluble crosslinking fraction and the pellet was further digested by a filter-aided sample preparation (FASP) digest. The FASP procedure was conducted in principle as described before[95] but with the following modifications: Samples were lysed in buffer containing 100 mM dithiothreitol (DTT), 4% (w/v) SDS and 100 mM HEPES (pH 7.3) by incubation at 75–80 *^◦^*C for 10 min. An 8 M urea buffer (prepared fresh in 100 mM HEPES, pH 7.3) was used for protein solubilization and filter-based cleanup to minimize urea-derived peptide modifications. For FASP, 30 µl of lysate was diluted with 200 µl urea buffer and loaded onto a centrifugal filter unit (Microcon-30, 30 kDa,Ultracel PL, Merck). Samples were centrifuged at 14,000g for 20 min at room temperature. This loading step was repeated until the entire sample was processed. Filters were washed twice with 200 µl urea buffer, followed by alkylation with 100 µl of 100 mM iodoacetamide (IAA) in 8 M urea buffer for 20 min at room temperature in the dark. Filters were subsequently washed twice with urea buffer and three times with 100 mM HEPES (pH 7.3), followed by centrifugation until near dryness. Proteins were digested on-filter with trypsin (500 ng per sample in 50 µl HEPES buffer) overnight at 37 *^◦^*C. Peptides were collected by centrifugation, followed by an additional elution step with 50 µl HEPES buffer. For secondary digestion, chymotrypsin (500 ng in 50 µl HEPES buffer) was added and incubated for 4 h at 37 *^◦^*C. Chymotrypsin digests were kept separately to the trypsin digest and not considered for further analysis. Trypsin digests were labelled thereafter as insoluble crosslink fraction and further processed by peptide size exclusion chromatography. Peptides were desalted using C18 cartridges (Sep-Pak C18, Waters). Columns were activated with methanol and equilibrated with 0.1% (v/v) TFA. Samples (adjusted to pH 3) were loaded, washed three times with 0.1% TFA, and eluted with 80% (v/v) acetonitrile (ACN) in 0.1% TFA. Eluates were dried in a vacuum concentrator and lyophilized for further peptide fractionation.

### 4.13 Crosslink peptide enrichment for tBuPhoX samples using FeNTA magnetic beads

tBuPhoX-crosslinked peptides were enriched after clean up using FeNTA magnetic beads (Cube Biotech). For 200 µg input material, 10 µl of bead slurry (25% stock; corresponding to 50 µl of 5% bead suspension) was used. Beads were equilibrated by washing three times with 1 ml of 80% (v/v) acetonitrile (ACN) in 0.1% (v/v) trifluoroacetic acid (TFA). Desalted peptide samples were resuspended in 80% ACN, 0.1% TFA and incubated with the equilibrated beads (final volume 500 µl) for 2 h at room temperature under continuous rotation. Beads were subsequently washed three times with 80% ACN, 0.1% TFA. Bound peptides were eluted twice with 50% (v/v) ACN, 2.5% (v/v) ammonium hydroxide for 1 min. Eluates were pooled and immediately neutralized with formic acid. Samples were dried in a vacuum concentrator and resuspended in 0.1% TFA prior to further analysis. Enriched tBuPhoX samples were subsequently subjected to size-exclusion chromatography.

### 4.14 Peptide size exclusion fractionation

Purified samples were reconstituted in 0.1% TFA to a final concentration of 3 ug/uL. 60 to 100 ug of peptides were injected per sample and condition on a Vanquish™ Analytical Purification LC System (Thermo Fisher Scientific) consisting of autosampler, SD-pumps, and UV detectors. Fractions have been collected automatically using an integrated Thermo Scientific™ Vanquish™ Fraction Collector. Peptides were separated on a TSKgel SuperSW2000 column (4.6 mm ID x 30 cm L, P/N: 0018674, Tosoh Bioscience) at a flow rate of 300 *µ*L min*^−^*^1^ using the SEC mobile phase (30% ACN in 0.1% TFA) and an isocratic gradient. The separation was monitored by UV absorption at 214 and 280 nm. Half-a-minute fractions (150 *µ*l) were collected into 0.6 uL low-bind reaction tubes over a separation window of 6 min. For analysis by liquid chromatography (LC)-MS/MS, fractions of interest (retention times 6-12 min) were collected and evaporated to dryness.

### 4.15 Mass spectrometry

### 4.16 Data-dependent acquisitions (DDA)

LC-MS/MS analysis was performed using an Orbitrap Eclipse (*C. elegans* nuclei crosslinking experiments) or Orbitrap Astral mass spectrometer with high-field asymmetric ion mobility spectrometry (FAIMS) interface (FAIMS Pro Duo, Thermo Fisher Scientific, Waltham, Massachusetts, United States) coupled with an EASY-Spray source and Vanquish Neo UHPLC system (Thermo Fisher Scientific). A trap column PepMap C18 (5 mm × 300 *µ*m ID, 5 *µ*m particles, 100 Å pore size) (Thermo Fisher Scientific, Waltham, Massachusetts, United States) and an analytical column Aurora Ultimate (250 mm × 75 *µ*m ID, 1.7 *µ*m, 120 Å)(Ion Opticks, Fitzroy, Australia) were employed for separation. The column temperature was set to 50°C. Sample loading was performed using 0.1% trifluoroacetic acid in water with a flow rate of 25 *µ*L min*^−^*^1^. Mobile phases used for separation were as follows: A 0.1% formic acid (FA) in water; B 80% acetonitrile, 0.1% FA in water. Peptides were eluted using a flow rate of 300 nL/min, with the following gradient: from 2% to 37% phase B in 60 min, from 37% to 47% phase B in 10 min, from 47% to 95% phase B in 1 min, followed by a washing step at 95% for 4 min, and re-equilibration of the column. The gradient was altered for the gradient optimization experiments to facilitate longer gradients (70 and 90 min gradient time). The mass spectrometry settings on the Astral mass spectrometer were set as follows: FAIMS separation was performed with the following settings: inner and outer electrode temperatures were 100°C, FAIMS carrier gas flow was 3.5 L/min, compensation voltages (CVs) of −48, −60, and −75 V were used in a stepwise mode during the analysis. The ion transfer tube temperature was set to 275°C. The mass spectrometer was operated in a data-dependent mode with cycle time 1s, using the following full scan parameters: m/z range 350-1300, nominal resolution of 180 000, with a target of 500% charges for the automated gain control (AGC), and a maximum injection time of 6 ms. For higher-energy collisional dissociation (HCD) MS/MS scans, single normalised collision energy (NCE) of 32% (non-cleavable crosslinker) or stepped HCD of 21, 25, 30% (cleavable crosslinker) was used. Precursor ions were isolated in a 1.6 Th (+/− 0.8 Th) window with no offset and accumulated for a maximum of 20 ms or until the AGC target of 500% was reached. Precursors of charge states from 3+ to 6+ were scheduled for fragmentation. Previously targeted precursors were dynamically excluded from fragmentation for 15 seconds. The sample load was typically in a range of 1 ng to 500 ng as indicated in the respective figure or with 250 ng for samples not part of the dilution experiment. Detailed parameters can be found in each raw file under the instrument method section. The mass spectrometry settings on the Eclipse mass spectrometer were set as follows: FAIMS separation was performed with the following settings: inner and outer electrode temperatures were 100°C, FAIMS carrier gas flow was 4.4 L/min, compensation voltages (CVs) of −50, −60, and −75 V were used in a stepwise mode during the analysis. The ion transfer tube temperature was set to 275°C. The mass spectrometer was operated in a data-dependent mode with cycle time 1s, using the following full scan parameters: m/z range 375-1300, nominal resolution of 120 000, with a target of 250% charges for the automated gain control (AGC), and a maximum injection time of 246 ms. For higher-energy collisional dissociation (HCD) MS/MS scans, single normalised collision energy (NCE) were set as for Astral measurements. Precursor ions were isolated in a 1.6 Th window with no offset and accumulated for a maximum of 70 ms or until the AGC target of 500% was reached. The resolution for MS2 scans was set to 30000. Precursors of charge states from 3+ to 6+ were scheduled for fragmentation. Previously targeted precursors were dynamically excluded from fragmentation for 15 seconds.

### 4.17 Data-independent acquisition (DIA) parameters

Data-independent acquisition (DIA) was performed on an Orbitrap Astral mass spectrometer operated in positive ion mode. The total method duration was either 70 or 90 min. FAIMS was employed in standard resolution mode with a single compensation voltage (CV) of −48 V or to test the impact of cycle time vs identification rate and data points per peak with combination −48 V, −60 V and −48 V, −60 V, −75 V. Full MS (MS1) scans were acquired in the Orbitrap at a resolution of 180,000 over an m/z range of 350–1200, with a maximum injection time of 6 ms and an absolute AGC target of 5 × 10^6^ ions. RF lens was set to 55%, and one microscan was acquired per scan. Advanced peak determination was enabled. DIA MS/MS scans were acquired using the Astral analyzer over an m/z range of 150–2000, with precursor isolation covering 350–1200 m/z. A total of 106 DIA windows were used with a fixed isolation width of 5, 8, 12, 16, 20 m/z and 0.5 m/z overlap or an variable window DIA method. MS2 spectra were acquired with a maximum injection time of 40 ms and an AGC target of 1 × 10^5^ ions (normalized AGC target 1000%). Fragmentation was performed using higher-energy collisional dissociation (HCD) with a normalized collision energy of 32%. Both MS1 and MS2 scans were acquired in centroid mode. Source fragmentation and lock mass correction were not applied. The acquisition scheme was optimized to balance cycle time and sampling density across chromatographic peaks. For DIA method evaluation the following parameters were tested for best performance with regard to maximise data points per elution peak and selectivity: MS1 injection time: 3, 6, 10, 30, 50, 100 ms; MS2 injection time: 5, 10, 20, 30, 40, 50, 60 ms; AGC MS2: 200, 400, 600, 800, 1000 ms. These parameters were altered for DIA 8 m/z, DIA 12 m/z and the variable window DIA method in order to find the optimal method.

### 4.18 Data analysis

Raw files for the Cas9 QC samples were analysed using Thermo Proteome Discoverer (v. 3.1.0.638). Searches were performed against the Cas9 sequence (Uniprot ID: Q99ZW2) plus a Crapome database (downloaded from https://www.thegpm.org/crap/). The workflow and search parameters were used as described in Müller at el. 2025[55]. In short, linear peptides were identified using MS Amanda search engine (v. 3.0.20.558)[96] and the crosslinked peptides were identified using MS Annika (v. 3.0)[64, 97, 98]. The workflow included per-file recalibration, followed by an initial search for linear peptides and monolinks. Spectra confidently assigned to linear peptides were excluded from subsequent analysis, and the remaining spectra were subjected to crosslink identification using MS Annika. False discovery rates (FDR) were estimated using the MS Annika validator node with a 1% threshold at both crosslink spectrum match (CSM) and residue pair levels, based on a target–decoy strategy. Automatic spectrum de-noising and scoring were applied as described previously[96]. *C. elegans* crosslinked nuclei data were analysed using xiSEARCH (v1.8.13)[99] and validated using xiFDR (v2.3.10)[53]. Crosslinked peptides were identified using xiSEARCH executed on a high-performance computing cluster (CBE) using a SLURM-based parallelization strategy. Searches were distributed across array jobs (n = the number of mgf files), with each task processing individual MGF files. Jobs were executed using Java (v11) with 8 CPUs and up to 350 GB RAM per task. Spectra were searched against a *C. elegans* nuclear protein database (3,295 entries, constructed from a linear search with the full *C. elegans* database (Uniprot Proteome UP000001940, 19804 entries, downloaded 05.09.2023) with a crosslinker-specific configuration for SDA, LC-SDA, BSP, DSSO, tBUPhoX. The config files are uploaded to PRIDE. Trypsin specificity (cleavage at K/R, up to two missed cleavages) was used, with a minimum peptide length of seven amino acids. Carbamidomethylation of cysteine was set as a fixed modification, while methionine oxidation and protein N-terminal acetylation were included as variable modifications. Crosslinker-derived modifications were restricted to lysine residues and peptide N-termini to reduce search space. All crosslinkers were defined as indicated in the respective config files and Table 2 with each crosslinker targeting lysine residues and protein N-termini. Precursor and fragment mass tolerances were set to 6 ppm and 15 ppm, respectively. Fragment ion types included b- and y-ions, and common neutral losses (H_2_O, NH_3_) were considered. Linear and mono-linked peptides were included in the search. Candidate selection was restricted to the top 100 alpha-peptides and top 5 alpha–beta combinations per spectrum to control computational complexity. Only the top-scoring match per spectrum was reported.

False discovery rate (FDR) estimation was performed using xiFDR based on a target–decoy strategy. Filtering was applied using a minimum peptide length of six residues, a minimum peptide coverage of 0.25, and a minimum delta score of 0.25. Unique peptide-spectrum matches (PSMs) were defined as the best-scoring match per peptide, crosslink site, charge state, and modification state. No restriction on link ambiguity was applied. FDR thresholds were controlled at multiple levels, including PSM, peptide pair, protein group, and protein group pair levels, with a target FDR of 5% at the protein group pair level. Residue pair-level FDR was not explicitly restricted during filtering. FDR estimation incorporated iterative boosting across multiple scoring features, including delta score, fragment count, peptide coverage, and identification counts, to improve separation of target and decoy distributions.

UAP56 open (−ATP, −RNA) and clamped (+ATP, +15U RNA) conformations (Figure 9A and Supplementary Figure S3H,I) were predicted using AlphaFold 3[85] via the AlphaFold web server. Models were generated from the UAP56 amino acid sequence (UniProt: Q13838), including the N-terminal 10×His-3C tag (MQLSHHHHHHHHHHSSGLEVLFQGP), using a diffusion-based deep learning framework that iteratively refines atomic coordinates and provides confidence estimates. Predicted structures were visualized in UCSF ChimeraX (v1.9)[100] and crosslinks were mapped using XMAS[101], a ChimeraX extension enabling integration of crosslinking mass spectrometry data. The conversion of crosslinks from xiSEARCH/xiFDR to the XMAS format was performed in pyXLMS[102]. XMAS was further used to calculate *Cα* - *Cα* distances and to visualize crosslinks on the structural models.

For data filtering and visualisation Python 3.10.16 was used with the following main packages: pandas (v 2.2.3)[103, 104], numpy (v 2.2.1)[103, 105], matplotlib (v 3.10.0)[106] (pyplot, venn (v 3.10.0)), seaborn (v 0.13.2)[107], scipy (V 1.15.0) and bioinfokit (v 2.1.3)[108, 109], biopython (v 1.84)[110].

### 4.19 Fasta file construction and data organisation

For identification of crosslinked peptides and linear/monolink peptides several databases needed to be constructed to perform data analysis. Experiment-specific FASTA databases were tailored to each sample type. For E.coli ribosome crosslink experiments, an Escherichia coli K12 reference proteome (Uniprot Proteome UP000000625, 4,403 entries, 25.11.2024) was used to identify respective crosslinks that have been used to construct the spectral library for Human vs E.coli entrapment datasets. The resulting E.coli spectral library consisted of 323 target precursors and 969 decoys (DD, DT, TD for each precursor). The respective Human ribosome (Uniprot Proteome UP000005640, 20554 entries, downloaded 12.12.2024) fasta for crosslink identification resulted in 1105 protein sequences with the respective spectral library of 58 target precursors and 174 decoy entries. Targeted analyses of Cas9 samples employed a reduced database comprising Cas9 and common contaminants (Crapome; 119 entries) which resulted in a spectral library of 4503 target precursors and 13509 decoy entries. For structural and crosslinking analyses of the UAP56 helicase, custom databases containing the respective target protein and background proteins were used (357 entries). The spectral library used for DIA analysis in Spectronaut contained 370 target precursors and 1110 decoys. For *C. elegans* nuclei crosslinking experiments *Caenorhabditis elegans* proteome databases were employed, including a full UniProt reference proteome (Uniprot Proteome UP000001940, 19804 entries, downloaded 05.09.2023) and a reduced, experiment-specific nuclear protein database (6749 entries, 3695 nucleus annotated) to optimize search space and sensitivity for crosslink searches. The resulting spectral library used for DIA entrapment experiments consisted of 849 target precursors (tBuPhoX dataset) and 2547 decoys. The BSP entrapment spectral library consisted of 323 target precursors and 969 decoys. We also employed a third *C. elegans* data set as entrapment dataset previously published with the PRIDE identifier PXD055488. All dependencies of fasta files to raw files and data analysis strategies are summarised in Summary Table 1.

**Table 1:** Special reagents used in this study.

| Reagent name | Catalogue number | Supplier |
| --- | --- | --- |
| Cas9 from *S. pyogenes* fused with a Halo-tag | In house | Deng et al.[91] |
| tBuPhoX (TBDSPP) | A52287 | Thermo Fisher Scientific |
| PhoX (DSPP) | A52286 | Thermo Fisher Scientific |
| DSSO | A33545 | Thermo Fisher Scientific |
| SDA | 26167 | Thermo Fisher Scientific |
| LCSDA | 26168 | Thermo Fisher Scientific |
| BSPNO2 | Literature, synthesised at Kandioller Group | Chowdhury et al.[57] |
| *E. coli* Ribosome | P0763S | New England Biolabs |

**Table 2:**
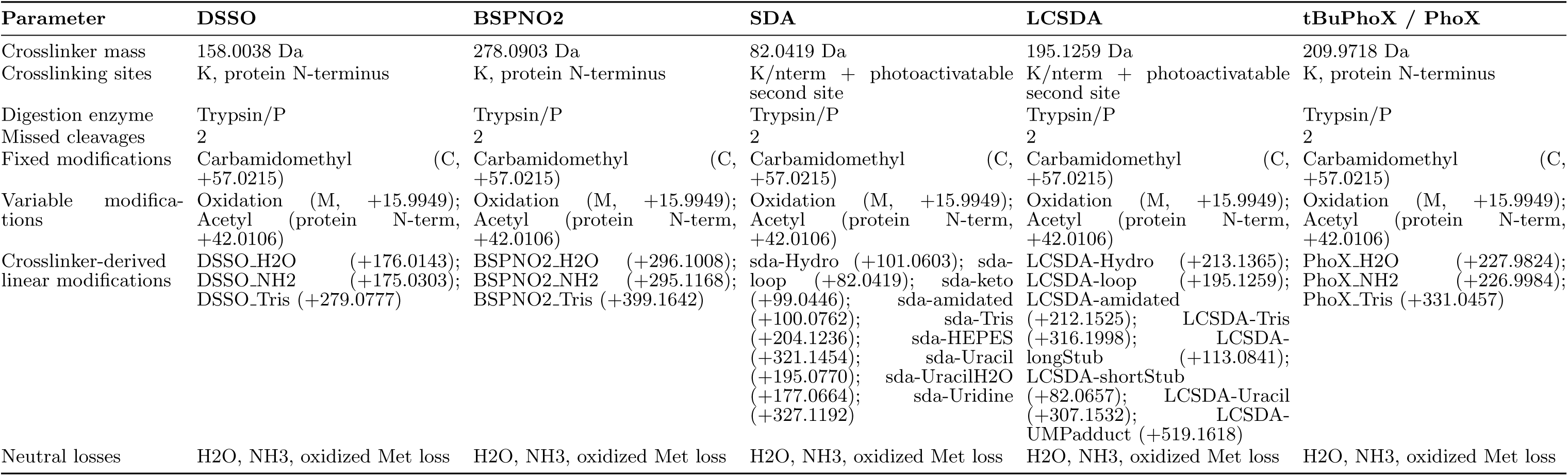
Crosslinker-specific search settings, modifications and crosslinker masses.

### 4.20 GO-term enrichment analysis

Gene Ontology (GO) enrichment analysis was performed using the g:Profiler Python interface (GProfiler). Independent enrichment analyses were conducted for each gene set using the *Caenorhabditis elegans* reference database and the Biological Process ontology (GO:BP). For each dataset, enriched GO terms were ranked based on adjusted P values, and the top-ranking terms were selected for downstream comparison. Overlapping GO terms between datasets were identified by matching GO identifiers, and enrichment significance was transformed to −*log*_10_ (P value) for visualization. Multiple testing correction was performed internally by g:Profiler using its default g:SCS procedure, and reported P values correspond to adjusted significance values. Comparative analysis focused either on the top enriched terms or, where indicated, on the subset of terms exhibiting the largest differences in enrichment significance between conditions, defined by the absolute difference in −*log*_10_ (P value).

### 4.21 Creating a Spectral library with Decoy Spectra

The proposed spectral library creation workflow is implemented in the programming language python as a standalone script with command line interface and requires at least three files as input: Mass spectra in either mzML or MGF format, crosslink search engine results containing crosslink-spectrum-matches (CSMs) either from MS Annika[64] or xiSearch[99] with xiFDR[53], and a configuration file with additional parameters.

The preferred input format for mass spectra is mzML as it contains additional information beneficial for spectral library creation compared to the MGF counterpart. We recommend converting Thermo RAW files using ThermoRawFileParser[111] to mzML format which preserves most of the spectral information and assures readable spectrum headers that are crucial for matching CSMs to their corresponding spectra. The supplied crosslink search engine results should already be validated at the CSM-level to make sure that only high-quality spectra and identifications are used for the spectral library. We suggest using CSMs that are below the 1% false-discovery-rate (FDR) threshold or stricter. The accepted input formats are Microsoft Excel (xlsx) format for MS Annika results and comma-separated values (csv) format for xiFDR results. Furthermore, the spectral library creation process is controlled by the configuration file that needs to be adopted by the user before running the python script. The configuration file contains information such as which crosslinker is used, which post-translational modifications (PTMs) are to be expected, ion types, fragment match tolerance, and other parameters that steer the annotation and spectral library creation workflow. For cleavable crosslinkers it is additionally required that all stubs are given as delta masses in the configuration file in order to properly annotate fragment ions containing crosslinker stubs. We provide an example configuration file together with the python script that users can freely adopt and that highlights the format and structure of the file. The required computational resources for running the python script are largely dependent on the size and amount of the mass spectrum files. While considerable effort went into optimizing the loading and annotation of spectra, we still need to load and store all spectra of a file into memory uncompressed to assure proper and timely matching to CSMs and therefore advise at least 32 GB of memory to ensure a smooth spectral library creation process. Spectra files are loaded using the python package pyteomics[112, 113] and we store precursor information, retention time, compensation voltage and the maximum observed intensity internally in addition to peak information. Crosslink search engine result files are parsed manually and CSMs are then matched to their corresponding mass spectra via their scan number. Scan numbers are extracted from mass spectra via the spectrum header by looking for an integer number that is prefixed by a “scan=” string. This correctly parses the scan number for mzML and MGF files created with ThermoRawFileParser[111] but can be adapted via the configuration file for other conversion tools.

For each CSM the script then calculates all possible theoretical fragment ion masses based on the identified crosslinked peptidoforms (peptides and their associated PTMs including crosslink modifications) also using pyteomics. Subsequently these theoretical ion masses are then matched to the experimental mass spectrum’s peaks given a user-defined tolerance (by default 0.02 Da) and non-annotated peaks are removed from the spectrum. This annotated and cleaned spectrum is added as a target (or in crosslink terms target-target, abbreviated as TT) spectrum to the spectral library. Furthermore, the next step in the workflow is the generation of plausible decoy spectra for every TT spectrum which differs significantly for crosslink spectral library creation compared to traditional linear peptide proteomics due a different target-decoy relationship. To rehearse, in crosslinking proteomics any match can be a combination of target and decoy peptides, essentially forming four distinct groups: target-target or TT matches as previously introduced, and additionally target-decoy (TD), decoy-target (DT), and decoy-decoy (DD) matches. To preserve this distinct behaviour, the script calculates three different decoy spectra based on the originally – in the previous step – annotated target-target spectral library spectrum. Specifically, decoys are generated using the reverse algorithm described by Zhang et al.[65]: Essentially, the target peptide is reversed except for the C-terminal residue and all PTMs stay attached to their corresponding amino acids (or the N- or C-terminus). The m/z values of all fragment ions are then shifted to correspond to the new decoy peptide sequence while conserving fragment ion intensities. For the TD and DT spectra this step is only performed for one of the respective peptides (either the beta- or alpha-peptide correspondingly) while DD spectra are generated by transforming both peptides to decoys. In total we end up with three decoy spectra per target spectrum in our spectral library. A visualization of the procedure is given in section “A four-state target–decoy spectral library strategy for DIA-based crosslinking mass spectrometry” in Figure 3. Finally, supplementary information on both spectrum-level (e.g. compensation voltage) and identification-level (e.g. associated proteins and crosslinked residues) are added to the spectral library. The spectral library is then exported in standard comma-separated values (csv) format that conforms with the input requirements of Spectronaut. In addition to the full spectral library, the script also exports the TT, TD, DT, and DD libraries separately which would allow users to adapt them or explore different decoy generation methods. Version 1.4.22 of the spectral library creation script was used for the analyses in this manuscript. The script, configuration file, example data, and documentation is freely available under a permissive MIT open source license via the GitHub repository https://github.com/hgb-bin-proteomics/MSAnnika_Spectral_Library_exporter.

We also want to acknowledge the previously published spectral library script (available via https://github.com/Rappsilber-Laboratory/xiDIA-library)[37] here. While our workflow/script is a complete standalone implementation with its own annotation and features, their work helped in understanding the spectral library creation process and the Spectronaut specific spectral library format.

### 4.22 DIA data analysis in Spectronaut

Data-independent acquisition (DIA) data were analyzed using Spectronaut (v20.4, Biognosys) in peptide-centric mode. A project-specific spectral library was used, containing 4,506 target precursors and 13,518 decoy entries to enable target–decoy-based scoring and false discovery rate (FDR) estimation. The spectral library differed depending on the experiment and sample analysed (Summary Table 1). Raw data were processed using the Spectronaut default pipeline, including automatic mass calibration, iRT-based retention time alignment, and gradient fine-structure correction. Ion chromatograms were extracted using dynamic m/z and retention time windows with maximum intensity extraction, followed by machine learning–based scoring of precursor identifications. Those settings were left on default. Precursor identifications were filtered at a q-value threshold of 0.01 (1% FDR) at both run and experiment levels. Protein-level FDR (here unique link FDR (major group)) was controlled at 1% (experiment) and 5% (run level). A kernel density estimator was used for p-value estimation. Duplicate assays were excluded, and in-source fragment filtering was set to retain “likely in-source fragmented” peptides. Decoy generation within Spectronaut was disabled, as decoys were pre-defined in the spectral library. Quantification was performed at MS1 and MS2 level using fragment ion peak areas (area-based quantification). Cross-run normalization was enabled using the automatic normalization strategy (global normalisation). Interference correction was enabled and restricted to identified peptides only, requiring a minimum of 2 MS1 and 3 MS2 fragment ions per precursor. Quantification was performed on modified peptide level (grouped by modified sequence), and TopN filtering was applied (1–10 precursors per peptide/protein group). A *log*_2_ intensity filter (minimum *log*_2_ intensity ≥ 0) was applied. No missing value imputation was performed. Protein-RNA crosslinks and monolinks were quantified using the PTM Candidate workflow (direct DIA workflow with automated library and decoy generation), treating crosslinks as modified peptide species. PTM localization was enabled. Quantification was based on MS2-level precursor areas aggregated across runs. Differential abundance between conditions was assessed using an unpaired two-sided t-test without assuming equal variance. Significance thresholds were defined using p-values (Benjamini–Hochberg correction) with a cutoff of p *<* 0.05 and Qvalue *<* 0.05. An additional *log*_2_ fold-change filter of 1x fold change was applied during candidate selection. Data completeness, recovery, and coefficient of variation (CV) were calculated within Spectronaut across all DIA conditions. Candidate tables were exported from Spectronaut for further analysis and are collected as Summary Tables 2. Representative spectra of protein-RNA links are shown in Supplementary Figure 10-12.

### 4.23 Quantitative crosslinking data processing and statistical analysis

DIA-derived peak area data were processed in Python using custom scripts. Normalized MS1 peak areas (retrieved from Spectronaut) were aggregated at the level of unique residue pairs (URPs) by grouping replicate measurements within each condition. For each URP, replicate-level means were calculated and subsequently summarized across replicates to obtain condition-level mean intensities. Crosslinked peptides were classified based on replicate completeness using a detection-based filtering strategy. Features quantified in ≥2 out of 3 replicates in both conditions were retained as high-confidence (“both conditions”), whereas features detected in only one condition were assigned as condition-specific (open or clamped). Features with *<*2 detected replicates in either condition were excluded from downstream analysis. Detection was defined based on positive signal intensity after normalization. For partially observed features (2/3 replicates), missing values were imputed to reconstruct complete triplicate profiles. Imputation was performed within each URP and condition only if ≥2 replicate measurements were available, and missing values were replaced by the mean of observed replicate intensities. This approach minimized bias while preserving quantitative variance structure. Quantitative profiles were visualized using *log*_10_-transformed normalized peak areas across replicates and conditions, retaining replicate-level resolution to enable direct comparison of abundance changes between conformational states. Representative spectra of a quantified crosslinked peptide in Spectronaut is shown in Supplementary Figure S9.

### 4.24 DIA data analysis in Skyline

Crosslinked peptides were quantified using Skyline (64-bit, v26.1.0.057 (c07debd50))[69, 70], which provides native support for crosslinking workflows. A spectral library was generated within Skyline using the peptide settings wizard (“Build” function), with default parameters. Library construction was based on an .ssl file containing crosslink identifications (including file name, scan number, charge state, peptide sequence, score type and score: 0.05), together with the corresponding .mgf files located in the same directory. Crosslinked peptide sequences were defined in the .ssl file using the format: peptide A–peptide B–[crosslinker@residue A,residue B] (e.g., SFRIFNDPKNV–VASIPRHNVLSNGHLMEKLQSNLE–[PhoX@4,5]).

For data analysis, Skyline peptide settings were configured with tryptic specificity (KR/P), allowing up to three missed cleavages, with peptide lengths ranging from 6 to 50 amino acids. Carbamidomethylation (C), methionine oxidation, and PhoX crosslinker modifications were included. Crosslinkers were defined as structural modifications with lysine specificity. All other parameters were kept at default settings. Transition settings included precursor charges of 3–6 and fragment ion charges of 1–2. Fragment ions comprised b-, y-, and precursor ions, spanning from the precursor to the last detectable ion. The top three product ions were selected from the spectral library. Instrument settings specified a product ion m/z range of 350–1200. Full-scan settings were configured for Orbitrap acquisition at 180,000 resolving power, with DIA-based MS/MS filtering using centroid data and a 12 m/z isolation window (0.5 m/z margin), and a mass accuracy of 10 ppm. Raw files acquired on the Orbitrap Astral instrument were imported into Skyline (File → Import → Results). Chromatographic peaks were extracted for each precursor[114], and quantitative data were exported using Skyline’s reporting tools (File → Export → Report). Skyline enabled visualization of crosslinked peptides as paired sequences with annotated crosslink sites, and fragment ions were represented using standard ion notation. Of note, at this moment it is not possible to include a decoy library specific for crosslinking data into Skyline, therefore exported data were filtered to either 0.7 or 0.95 idotp value to increase the confidence in the crosslinking data.

### 4.25 Rescoring of quantitative DIA crosslinking data

The proposed rescoring workflow consists of two steps: first the calculation of new features and scores that should be used for rescoring, and second the rescoring step which is performed with Mokapot [80]. The first step is implemented as a postprocessing script in python which takes the result from Spectronaut and the spectral library as input. Among other features such as the number of matched ions per peptide, the sequence coverage per peptide, and the number of fragments containing crosslink modifications, we also calculate the following scores:

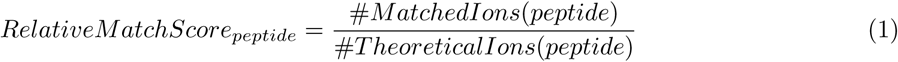

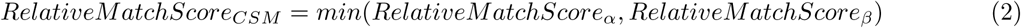

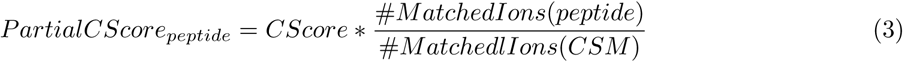

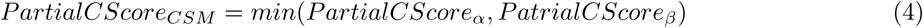

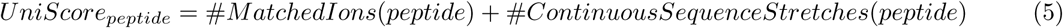

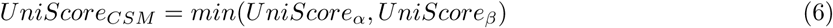

The RelativeMatchScore (equation 1) is a very basic measurement of how many fragment ions were matched in relation to the total number of theoretically possible fragment ions. The CScore is the score returned by Spectronaut itself and the UniScore (equation 5) we calculate as proposed by Tabata et al. [79]. The PartialCScore (equation 3) for a peptide is simply the CScore multiplied with the fraction of fragment ions of the peptide in relation to the total number of matched ions for the corresponding CSM. The CSM scores (equations 2, 4, 6) are in all cases the minimum score of both (*α* and *β*) crosslinked peptides which is an approach that we also use in the crosslink search engine we developed and which has proven to be effective for FDR estimation [64]. Finally, the postprocessing script creates a new table with the new features and scores to be used with Mokapot. Additionally a table for direct usage and validation with xiFDR [53] is created (without machine learning-based rescoring). The second step is implemented as a separate python script which takes the output of the postprocessing step as input. We implemented and explored two different rescoring approaches: a) direct rescoring of a CSM by using features associated with the entire CSM (denoted as CSM-level rescoring), and b) rescoring of the two individual crosslinked peptides of the CSM by splitting the CSM up into two PSMs, rescoring the two PSMs separately based on features associated with the PSM, and then calculating the CSM score as the minimum of the two rescored PSM scores (denoted as PSM-level rescoring). The rescoring is performed with Mokapot (version https://github.com/wfondrie/mokapot, main@2b1f13a, most recent version as of April 2026) using the Percolator [81] model emulation. The rescoring script outputs the original data with an extra “Mokapot Score” column which includes the new learned score, which then can be used for validation and FDR estimation. For assessing the rescoring we did target-decoy based FDR estimation using the formula shown in equation 7 and analysing the number of entrapment hits as well as calculation of an empirical FDR based on the entrapment. The empirical FDR is calculated by simply dividing the number of entrapment hits (denoted as E in figures) by the number of non-entrapment hits (target system, denoted as T in figures).

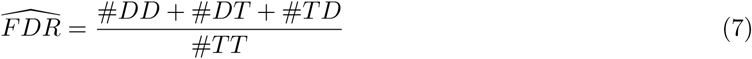

For aggregation to unique residue pairs and FDR estimation we used the python package pyXLMS[102] (version 1.7.5) with standard parameters and a target FDR of 1%. We also provide a parser for both rescored and non-rescored results for direct usage with pyXLMS. For this manuscript version 1.2.10 of the postprocessing script and version 1.1.4 of the rescoring script were used. The complete source code, an extensive list of all calculated features, and the figure code are given in the project’s GitHub repository and the rescoring submodule accessible via https://github.com/hgb-bin-proteomics/MSAnnika_Spectral_Library_exporter.

## 5 Data availability

The mass spectrometry proteomics data have been deposited to the ProteomeX-change Consortium (http://proteomecentral.proteomexchange.org) via the PRIDE partner repository[115, 116] with the dataset identifier PXD079320. The C.elegans protein-protein interaction community dataset was deposited separately with the identifier PXD079460. This study also employed published data with the identifier PXD059096[55] and identifier PXD055488[64]. Source data are provided with this paper and includes raw tables from Spectronaut analysis (directDIA, crosslink quantitation, entrapment experiments), xiFDR crosslink tables (CSM, Link), Candidate tables for protein-RNA links and Monolinks, Supplementary Excel tables and Summary tables.

## 6 Code Availability

All custom code including scripts for spectral library creation, postprocessing, and rescoring are available under a permissive MIT open-source license via the GitHub repository https://github.com/hgb-bin-proteomics/MSAnnika_Spectral_Library_exporter which includes the rescoring code as a submodule. The rescoring code can also be separately accessed via https://github.com/hgb-bin-proteomics/MSAnnika_Spectral_Library_exporter_rescoring. The spectral library creation and post processing scripts require at least python version 3.7 to run but we recommend python 3.14 for its performance improvements. The rescoring script requires at least python version 3.12 and should be used with the pinned package versions as given in the repository. We recommend at least 32 GB of memory for running the spectral library creation script to ensure smooth and timely execution, especially for large input files.

## 7 Acknowledgements

We thank our colleagues at the Proteomics TechHub and Proteomics Core Facility for support and discussions, especially Julia Bubis, Karel Stejskal, Michael Schutzbier, Gabriela Krssakova, and Elisabeth Roithinger; the Bioinformatics Research Group for their support, especially Sebastian Dorl for his input on spectral libraries; the staff at the CLIP cluster (http://clip.science) for computational support; Andras Aszodi for help with setting up batch scripts to run xiSEARCH on the cluster; staff at the in-house Molecular Biology Services for aliquots of Cas9; Andrea Graziadei and Swantje Lenz for discussions on crosslink FDR and entrapment strategies; Ulrich Hohmann and Clemens Plaschka for discussions and suggestions regarding UAP56 RNA interactions and conformational changes. We further thank Ulrich Hohmann for providing UAP56, 15U RNA, and ATP aliquots, and Max Graf for help with protein SEC experiments. We also thank Thomas Lloyd Williams from the Electron Microscopy Facility for providing human ribosome aliquots. We further thank Manuel Matzinger for establishing the connection to the Kandioller Group and Angela Graf for worm collections. This work was funded by the ESPRIT programme, project number ESP566 (Grant DOI: 10.55776/ESP566, F.M.), the SFB projects F88-01-B and PAT2512023 (Grant DOIs: 10.55776/F88 and 10.55776/PAT2512023, F.M. and S.S.G.), and project P35045-B (Grant DOI: 10.55776/P35045, M.J.B. and V.D.), as well as project PAT2059025 (Grant DOI: 10.55776/PAT2059025, M.J.B. and V.D.) of the Austrian Science Fund (FWF). We thank the University of Vienna for financial support of the Kandioller Group at the Institute of Inorganic Chemistry, which enabled funding of P.G. and W.K. All LC–MS/MS analyses in Vienna were performed using the Vienna BioCenter Core Facilities instrument pool. We thank the Proteomics Core Facility, headed by Elisabeth Roitinger, for support and assistance with setting up the Orbitrap Eclipse and Orbitrap Astral mass spectrometers. Crosslinker characterization experiments were performed at the VBCF Metabolomics Facility. The Vienna BioCenter Core Facilities (VBCF) Metabolomics Facility acknowledges funding from the Austrian Federal Ministry of Education, Science and Research and the City of Vienna. DH is a member of the Max Perutz Labs (MPL) Mass Spectrometry Facility, we want to thank the MPL Mass Spectrometry Facility for their support and helpful discussions. This research was funded in whole, or in part, by the Austrian Science Fund (FWF). For the purpose of open-access publication, the authors have applied a CC BY public copyright licence to any Author Accepted Manuscript version arising from this submission. Large language models were used to assist with language editing and improving the readability of the manuscript.

## 8 Author contributions

MJB implemented the spectral library generation script, post-processing and rescoring script, further developed the idea of rescoring crosslinks, and performed data analysis and visualization (figures, structural models, plots) for Figure 3, 8, writing – original draft together with FM, writing – review & editing together with FM; SSG performed *C. elegans* cultures, worm collection and nuclei isolation for all *C. elegans* related experiments, wrote the corresponding method sections; PG synthesised the BSPNO2 crosslinker and wrote the corresponding method sections; GG verification of the BSPNO2 crosslinker by mass spectrometry and kinetic study of the stability of the crosslinker; DH provided the original idea and implementation of rescoring of crosslinked peptides using Mokapot; WK supervised PG and ensured financing of the BSPNO2 sub project; KM discussed the data during group meetings; VJ supervised SSG and acquired funding; VD supervised MJB and acquired funding; FM supervised and conceptualized the study, had the original idea of DIA crosslinking (DIA-QCLMS), designed and performed proteomics sample processing as well as MS experiments, purified the BSPNO2 crosslinker, performed data analysis and visualization (figures, structural models, plots) for Figure 1, 2, 4, 5, 6, 7, 9, writing – original draft together with MJB, writing – review & editing together with MJB, FM co-supervised MJB and SSG, funding acquisition (FWF ESPRIT ESP566); VJ, VD and FM supervised the study. All authors revised and agreed on the manuscript.

## 10 Supplementary information

The online version contains supplementary material. The supplement will be available after Peer Review.

## References

[1] Dill, K.A., MacCallum, J.L.: The Protein-Folding problem, 50 years on. Science (2012) 10.1126/science.1219021

[2] Alberts, B.: The cell as a collection of protein machines: Preparing the next generation of molecular biologists. Cell 92(3), 291–294 (1998) 10.1016/S0092-8674(00)80922-8

[3] Henzler-Wildman, K., Kern, D.: Dynamic personalities of proteins. Nature 450(7172), 964–972 (2007) 10.1038/nature06522

[4] Vendruscolo, M., Fuxreiter, M.: Protein condensation diseases: therapeutic opportunities. Nature Communications 13(1), 5550 (2022) 10.1038/s41467-022-32940-7

[5] Motlagh, H.N., Wrabl, J.O., Li, J., Hilser, V.J.: The ensemble nature of allostery. Nature 508(7496), 331–339 (2014) 10.1038/nature13001

[6] Changeux, J.-P., Christopoulos, A.: Allosteric modulation as a unifying mechanism for receptor function and regulation. Diabetes, Obesity and Metabolism 19, 4–21 (2017) 10.1016/j.cell.2016.08.015

[7] Deribe, Y.L., Pawson, T., Dikic, I.: Post-translational modifications in signal integration. Nature Structural & Molecular Biology 17(6), 666–672 (2010) 10.1038/nsmb.1842

[8] Nooren, I.M.A., Thornton, J.M.: Structural characterisation and functional significance of transient protein–protein interactions. Journal of Molecular Biology 325(5), 991–1018 (2003) 10.1016/S0022-2836(02)01281-0

[9] Marsh, J.A., Teichmann, S.A.: Structure, dynamics, assembly, and evolution of protein complexes. Annual Review of Biochemistry 84, 551–575 (2015) 10.1146/annurev-biochem-060614-034142

[10] Yi, W., Yan, J.: Decoding RNA–Protein interactions: Methodological advances and emerging challenges. Advanced Genetics 6(2), 2500011 (2025) 10.1002/ggn2.202500011

[11] Ramanathan, M., Porter, D.F., Khavari, P.A.: Methods to study RNA–protein interactions. Nature Methods 16(3), 225–234 (2019) 10.1038/s41592-019-0330-1

[12] Aebersold, R., Mann, M.: Mass-spectrometric exploration of proteome structure and function. Nature 537(7620), 347–355 (2016) 10.1038/nature19949

[13] Wang, B., Xie, Z.-R., Chen, J., Wu, Y.: Integrating structural information to study the dynamics of protein-protein interactions in cells. Structure 26(10), 1414–14243 (2018) 10.1016/j.str.2018.07.010

[14] Speer, S.L., Zheng, W., Jiang, X., Chu, I.-T., Guseman, A.J., Liu, M., Pielak, G.J., Li, C.: The intracellular environment affects protein–protein interactions. Proceedings of the National Academy of Sciences 118(11), 2019918118 (2021) 10.1073/pnas.2019918118

[15] Carter, B., Justin, H.S., Gulick, D., Gamsby, J.J.: The molecular clock and neurodegenerative disease: A stressful time. Front. Mol. Biosci. 8, 644747 (2021) 10.3389/fmolb.2021.644747

[16] Wang, X., Wang, R., Li, J.: Influence of sleep disruption on protein accumulation in neurodegenerative diseases. Ageing and Neurodegenerative Diseases 2(1) (2022) 10.20517/and.2021.10

[17] Miller, M.D., Phillips, J., George N.: Moving beyond static snapshots: Protein dynamics and the protein data bank. Journal of Biological Chemistry 296, 100749 (2021) 10.1016/j.jbc.2021.100749

[18] Leone, V., Marinelli, F.: From snapshots to ensembles: Integrating experimental data and dynamics. Current Opinion in Structural Biology 95, 103155 (2025) 10.1016/j.sbi.2025.103155

[19] Ghadie, M.A., Xia, Y.: Are transient protein-protein interactions more dispensable? PLOS Computational Biology 18(4), 1010013 (2022) 10.1371/journal.pcbi.1010013

[20] O’Reilly, F.J., Rappsilber, J.: Cross-linking mass spectrometry: methods and applications in structural, molecular and systems biology. Nature Structural & Molecular Biology 25(11), 1000–1008 (2018) 10.1038/s41594-018-0147-0

[21] Piersimoni, L., Kastritis, P.L., Arlt, C., Sinz, A.: Cross-Linking mass spectrometry for investigating protein conformations and Protein–Protein InteractionsA method for all seasons. Chemical Reviews (2021) 10.1021/acs.chemrev.1c00786

[22] Yu, C., Huang, L.: New advances in cross-linking mass spectrometry toward structural systems biology. Current Opinion in Chemical Biology 76, 102357 (2023) 10.1016/j.cbpa.2023.102357

[23] Graziadei, A., Rappsilber, J.: Leveraging crosslinking mass spectrometry in structural and cell biology. Structure 30(1), 37–54 (2022) 10.1016/j.str.2021.11.007

[24] Liu, F., Rijkers, D.T.S., Heck, A.J.R.: Proteome-wide profiling of protein assemblies by cross-linking mass spectrometry. Nature Methods 12(12), 1179–1184 (2015) 10.1038/nmeth.3603

[25] James, E.I., Murphree, T.A., Vorauer, C., Engen, J.R., Guttman, M.: Advances in Hydrogen/Deuterium exchange mass spectrometry and the pursuit of challenging biological systems. Chemical Reviews (2021) 10.1021/acs.chemrev.1c00279

[26] Masson, G.R., Burke, J.E., Ahn, N.G., Anand, G.S., Borchers, C., Brier, S., Bou-Assaf, G.M., Engen, J.R., Englander, S.W., Faber, J., Garlish, R., Griffin, P.R., Gross, M.L., Guttman, M., Hamuro, Y., Heck, A.J.R., Houde, D., Iacob, R.E., Jørgensen, T.J.D., Kaltashov, I.A., Klinman, J.P., Konermann, L., Man, P., Mayne, L., Pascal, B.D., Reichmann, D., Skehel, M., Snijder, J., Strutzenberg, T.S., Underbakke, E.S., Wagner, C., Wales, T.E., Walters, B.T., Weis, D.D., Wilson, D.J., Wintrode, P.L., Zhang, Z., Zheng, J., Schriemer, D.C., Rand, K.D.: Recommendations for performing, interpreting and reporting hydrogen deuterium exchange mass spectrometry (HDX-MS) experiments. Nature Methods 16(7), 595–602 (2019) 10.1038/s41592-019-0459-y

[27] Malinovska, L., Cappelletti, V., Kohler, D., Piazza, I., Tsai, T.-H., Pepelnjak, M., Stalder, P., Dorig, C., Sesterhenn, F., Elsässer, F., Kralickova, L., Beaton, N., Reiter, L., Souza, N., Vitek, O., Picotti, P.: Proteome-wide structural changes measured with limited proteolysis-mass spectrometry: an advanced protocol for high-throughput applications. Nature Protocols 18(3), 659–682 (2022) 10.1038/s41596-022-00771-x

[28] Chen, Z.A., Rappsilber, J.: Quantitative cross-linking/mass spectrometry to elucidate structural changes in proteins and their complexes. Nat. Protoc. 14(1), 171–201 (2019) 10.1038/s41596-018-0089-3

[29] Keller, A., Bakhtina, A., Bruce, J.E.: Large-Scale quantitative Cross-Linking and mass spectrometry provide new insight into protein conformational plasticity within organelles, cells, and tissues. Journal of Proteome Research (2025) 10.1021/acs.jproteome.4c01030

[30] Luo, J., Ranish, J.: Isobaric crosslinking mass spectrometry technology for studying conformational and structural changes in proteins and complexes. eLife 13, 99809 (2024) 10.7554/eLife.99809

[31] Müller, F., Graziadei, A., Rappsilber, J.: Quantitative photo-crosslinking mass spectrometry revealing protein structure response to environmental changes. Analytical Chemistry 91(14), 9041–9048 (2019) 10.1021/acs.analchem.9b01339

[32] Jiang, T., Zhang, H., Da Silva, G.M., Gyawali, Y.P., Feng, C.: Deciphering mutational effects on inducible no synthase conformational dynamics via quantitative cross-linking mass spectrometry and alphafold2 subsampling. Journal of Biological Chemistry 301(11), 110673 (2025) 10.1016/j.jbc.2025.110673

[33] Jiang, T., Wan, G., Zhang, H., Gyawali, Y.P., Underbakke, E.S., Feng, C.: Mapping the intersubunit interdomain FMN-Heme interactions in neuronal nitric oxide synthase by targeted quantitative Cross-Linking mass spectrometry. Biochemistry (2024) 10.1021/acs.biochem.4c00157

[34] Mohammadi, A., Deroo, S., Leitner, A., Stengel, F., Krammer, E.-M., Aebersold, R., Prevost, M., Raussens, V.: Characterization of the n- and c-terminal domain interface of the three main apoe isoforms: A combined quantitative cross-linking mass spectrometry and molecular modeling study. Biochimica et Biophysica Acta (BBA) - General Subjects 1869(4), 130768 (2025) 10.1016/j.bbagen.2025.130768

[35] Mathay, M., Keller, A., Bruce, J.E.: Studying Protein-Ligand interactions by protein denaturation and quantitative Cross-Linking mass spectrometry. Anal Chem 95(25), 9432–9436 (2023) 10.1021/acs.analchem.2c04501

[36] Müller, F., Fischer, L., Chen, Z.A., Auchynnikava, T., Rappsilber, J.: On the reproducibility of Label-Free quantitative Cross-Linking/Mass spectrometry. Journal of The American Society for Mass Spectrometry (2017) 10.1007/s13361-017-1837-2

[37] Müller, F., Kolbowski, L., Bernhardt, O.M., Reiter, L., Rappsilber, J.: Data-independent acquisition improves quantitative cross-linking mass spectrometry. Molecular & Cellular Proteomics 18(4), (2019) 10.1074/mcp.TIR118.001276

[38] Müller, F., Rappsilber, J.: A protocol for studying structural dynamics of proteins by quantitative crosslinking mass spectrometry and data-independent acquisition. J Proteomics 218, 103721 (2020) 10.1016/j.jprot.2020.103721

[39] Rojas Echeverri, J.C., Hause, F., Iacobucci, C., Ihling, C.H., Tänzler, D., Shulman, N., Riffle, M., MacLean, B.X., Sinz, A.: A workflow for improved analysis of cross-linking mass spectrometry data integrating parallel accumulation-serial fragmentation with merox and skyline. Analytical Chemistry 96(19), 7373–7379 (2024) 10.1021/acs.analchem.4c00829

[40] Hao, Y., Chen, M., Huang, X., Xu, H., Wu, P., Chen, S.: 4D-diaXLMS: Proteome-wide Four-Dimensional Data-Independent acquisition workflow for Cross-Linking mass spectrometry. Analytical Chemistry (2023) 10.1021/acs.analchem.3c02824

[41] Belsom, A., Rappsilber, J.: Anatomy of a crosslinker. Current Opinion in Chemical Biology 60, 39–46 (2021) 10.1016/j.cbpa.2020.07.008

[42] Ziemianowicz, D.S., Ng, D., Schryvers, A.B., Schriemer, D.C.: Photo-cross-linking mass spectrometry and integrative modeling enables rapid screening of antigen interactions involving bacterial transferrin receptors. J. Proteome Res. 18(3), 934–946 (2019) 10.1021/acs.jproteome.8b00629

[43] Jiang, Y., Zhang, X., Nie, H., Fan, J., Di, S., Fu, H., Zhang, X., Wang, L., Tang, C.: Dissecting diazirine photo-reaction mechanism for protein residue-specific cross-linking and distance mapping. Nature Communications 15(1), 6060 (2024) 10.1038/s41467-024-50315-y

[44] Ziemianowicz, D.S., Ng, D., Schryvers, A.B., Schriemer, D.C.: Photo-cross-linking mass spectrometry and integrative modeling enables rapid screening of antigen interactions involving bacterial transferrin receptors. Journal of Proteome Research 18(3), 934–946 (2019) 10.1021/acs.jproteome.8b00629

[45] Pham, N.D., Parker, R.B., Kohler, J.J.: Photocrosslinking approaches to interactome mapping. Current Opinion in Chemical Biology 17(1), 90–101 (2013) 10.1016/j.cbpa.2012.10.034

[46] Ciancone, A., O’Reilly, F.J.: Photo-crosslinkers boost structural information from crosslinking mass spectrometry. Current Opinion in Structural Biology 93, 103102 (2025) 10.1016/j.sbi.2025.103102

[47] Taniguchi, I., Ohno, M.: ATP-dependent recruitment of export factor Aly/REF onto intronless mRNAs by RNA helicase UAP56. Mol Cell Biol 28(2), 601–608 (2008) 10.1128/MCB.01341-07

[48] Yamazaki, T., Fujiwara, N., Yukinaga, H., Ebisuya, M., Shiki, T., Kurihara, T., Kioka, N., Kambe, T., Nagao, M., Nishida, E., Masuda, S.: The closely related RNA helicases, UAP56 and URH49, preferentially form distinct mRNA export machineries and coordinately regulate mitotic progression. Mol Biol Cell 21(16), 2953–2965 (2010) 10.1091/mbc.e09-10-0913

[49] Pühringer, T., Hohmann, U., Fin, L., Pacheco-Fiallos, B., Schellhaas, U., Brennecke, J., Plaschka, C.: Structure of the human core transcription-export complex reveals a hub for multivalent interactions. Elife 9 (2020) 10.7554/eLife.61503

[50] Hohmann, U., Graf, M., Tirian, L., Pacheco-Fiallos, B., Schellhaas, U., Fin, L., Handler, D., Phillips, A.W., Riabov-Bassat, D., Faraway, R., Pühringer, T., Szalay, M.-F., Roitinger, E., Brennecke, J., Plaschka, C.: An ATP-gated molecular switch orchestrates human mRNA export. Nature 649(8098), 1042–1050 (2025) 10.1038/s41586-025-09832-z

[51] Xie, Y., Gao, S., Zhang, K., Bhat, P., Clarke, B.P., Batten, K., Mei, M., Gazzara, M., Shay, J.W., Lynch, K.W., Angelos, A.E., Hill, P.S., Ivey, A.L., Fontoura, B.M.A., Ren, Y.: Structural basis for high-order complex of SARNP and DDX39B to facilitate mRNP assembly. Cell Rep 42(8), 112988 (2023) 10.1016/j.celrep.2023.112988

[52] Clarke, B.P., Gao, S., Mei, M., Xie, D., Angelos, A.E., Vazhavilla, A., Hill, P.S., Cagatay, T., Batten, K., Shay, J.W., Xie, Y., Fontoura, B.M.A., Ren, Y.: Structural mechanism of ddx39b regulation by human trex-2 and a related complex in mrnp remodeling. Nature Communications 16(1), 5471 (2025) 10.1038/s41467-025-60547-1

[53] Fischer, L., Rappsilber, J.: Quirks of error estimation in Cross-Linking/Mass spectrometry (2017) 10.1021/acs.analchem.6b03745

[54] Bekker-Jensen, D.B., Bernhardt, O.M., Hogrebe, A., Martinez-Val, A., Verbeke, L., Gandhi, T., Kelstrup, C.D., Reiter, L., Olsen, J.V.: Rapid and site-specific deep phosphoproteome profiling by data-independent acquisition without the need for spectral libraries. Nat Commun 11(1), 787 (2020) 10.1038/s41467-020-14609-1

[55] Müller, F., Birklbauer, M.J., Bubis, J., Stejskal, K., Dorfer, V., Mechtler, K.: Breaking barriers in crosslinking mass spectrometry with enhanced throughput and sensitivity using orbitrap astral. Nat Commun 16(1), 9877 (2025) 10.1038/s41467-025-64844-7

[56] Escher, C., Reiter, L., MacLean, B., Ossola, R., Herzog, F., Chilton, J., MacCoss, M.J., Rinner, O.: Using iRT, a normalized retention time for more targeted measurement of peptides. PROTEOMICS 12(8), 1111–1121 (2012) 10.1002/pmic.201100463

[57] Chowdhury, S.M., Du, X., Tolic, N., Wu, S., Moore, R.J., Mayer, M.U., Smith, R.D., Adkins, J.N.: Identification of cross-linked peptides after click-based enrichment using sequential collision-induced dissociation and electron transfer dissociation tandem mass spectrometry. Anal Chem 81(13), 5524–5532 (2009) 10.1021/ac900853k

[58] Gao, H., Zhao, L., Zhong, B., Zhang, B., Gong, Z., Zhao, B., Liu, Y., Zhao, Q., Zhang, L., Zhang, Y.: In-Depth crosslinking in minutes by a compact, Membrane-Permeable, and Alkynyl-Enrichable crosslinker. Anal Chem 94(21), 7551–7558 (2022) 10.1021/acs.analchem.2c00335

[59] Zhao, L., An, Y., Zhao, N., Gao, H., Zhang, W., Gong, Z., Liu, X., Zhao, B., Liang, Z., Tang, C., Zhang, L., Zhang, Y., Zhao, Q.: Spatially resolved profiling of protein conformation and interactions by biocompatible chemical cross-linking in living cells. Nat Commun 15(1), 8331 (2024) 10.1038/s41467-024-52558-1

[60] Ren, Y., Schmiege, P., Blobel, G.: Structural and biochemical analyses of the DEAD-box ATPase sub2 in association with THO or yra1. Elife 6 (2017) 10.7554/eLife.20070

[61] Schuller, S.K., Schuller, J.M., Prabu, J.R., Baumgärtner, M., Bonneau, F., Basquin, J., Conti, E.: Structural insights into the nucleic acid remodeling mechanisms of the yeast THO-Sub2 complex. Elife 9 (2020) 10.7554/eLife.61467

[62] Pacheco-Fiallos, B., Vorländer, M.K., Riabov-Bassat, D., Fin, L., O’Reilly, F.J., Ayala, F.I., Schellhaas, U., Rappsilber, J., Plaschka, C.: mRNA recognition and packaging by the human transcription-export complex. Nature 616(7958), 828–835 (2023) 10.1038/s41586-023-05904-0

[63] Gromadzka, A.M., Steckelberg, A.-L., Singh, K.K., Hofmann, K., Gehring, N.H.: A short conserved motif in alyref directs cap- and ejc-dependent assembly of export complexes on spliced mrnas. Nucleic Acids Research 44(5), 2348–2361 (2016) 10.1093/nar/gkw009 https://academic.oup.com/nar/article-pdf/44/5/2348/17438108/gkw009.pdf

[64] Birklbauer, M.J., Müller, F., Geetha, S.S., Matzinger, M., Mechtler, K., Dorfer, V.: Proteome-wide non-cleavable crosslink identification with MS annika 3.0 reveals the structure of the c. elegans box C/D complex. Commun Chem 7(1), 300 (2024) 10.1038/s42004-024-01386-x

[65] Zhang, Z., Burke, M., Mirokhin, Y.A., Tchekhovskoi, D.V., Markey, S.P., Yu, W., Chaerkady, R., Hess, S., Stein, S.E.: Reverse and random decoy methods for false discovery rate estimation in high mass accuracy peptide spectral library searches (2018) 10.1021/acs.jproteome.7b00614

[66] Lancaster, N.M., Sinitcyn, P., Forny, P., Peters-Clarke, T.M., Fecher, C., Smith, A.J., Shishkova, E., Arrey, T.N., Pashkova, A., Robinson, M.L., Arp, N., Fan, J., Hansen, J., Galmozzi, A., Serrano, L.R., Rojas, J., Gasch, A.P., Westphall, M.S., Stewart, H., Hock, C., Damoc, E., Pagliarini, D.J., Zabrouskov, V., Coon, J.J.: Fast and deep phosphoproteome analysis with the orbitrap astral mass spectrometer. Nature Communications 15(1), 7016 (2024) 10.1038/s41467-024-51274-0

[67] Liu, H., Sadygov, R.G., Yates, J.R. 3rd: A model for random sampling and estimation of relative protein abundance in shotgun proteomics. Anal. Chem. 76(14), 4193–4201 (2004) 10.1021/ac0498563

[68] Kalxdorf, M., Müller, T., Stegle, O., Krijgsveld, J.: Icer improves proteome coverage and data completeness in global and single-cell proteomics. Nature Communications 12(1), 4787 (2021) 10.1038/s41467-021-25077-6

[69] MacLean, B., Tomazela, D.M., Shulman, N., Chambers, M., Finney, G.L., Frewen, B., Kern, R., Tabb, D.L., Liebler, D.C., MacCoss, M.J.: Skyline: an open source document editor for creating and analyzing targeted proteomics experiments. Bioinformatics 26(7), 966–968 (2010) 10.1093/bioinformatics/btq054

[70] Pino, L.K., Searle, B.C., Bollinger, J.G., Nunn, B., MacLean, B., MacCoss, M.J.: The skyline ecosystem: Informatics for quantitative mass spectrometry proteomics. Mass Spectrom Rev 39(3), 229–244 (2020) 10.1002/mas.21540

[71] Chavez, J.D., Eng, J.K., Schweppe, D.K., Cilia, M., Rivera, K., Zhong, X., Wu, X., Allen, T., Khurgel, M., Kumar, A., Lampropoulos, A., Larsson, M., Maity, S., Morozov, Y., Pathmasiri, W., Perez-Neut, M., Pineyro-Ruiz, C., Polina, E., Post, S., Rider, M., Tokmina-Roszyk, D., Tyson, K., Vieira Parrine Sant’Ana, D., Bruce, J.E.: A general method for targeted quantitative cross-linking mass spectrometry. PLoS One 11(12), 0167547 (2016) 10.1371/journal.pone.0167547

[72] Gupta, N., Pevzner, P.A.: False discovery rates of protein identifications: a strike against the two-peptide rule. J Proteome Res 8(9), 4173–4181 (2009) 10.1021/pr9004794

[73] Ma, J., Zhang, J., Wu, S., Li, D., Zhu, Y., He, F.: Improving the sensitivity of MASCOT search results validation by combining new features with bayesian nonparametric model. Proteomics 10(23), 4293–4300 (2010) 10.1002/pmic.200900668

[74] Granholm, V., Noble, W.S., Käll, L.: On using samples of known protein content to assess the statistical calibration of scores assigned to peptide-spectrum matches in shotgun proteomics. J Proteome Res 10(5), 2671–2678 (2011) 10.1021/pr1012619

[75] Müller, F., Brutiu, B.R., Saridakis, I., Leischner, T., Birklbauer, M.J., Matzinger, M., Madalinski, M., Lendl, T., Shaaban, S., Dorfer, V., Maulide, N., Mechtler, K.: Developing a new cleavable crosslinker reagent for in-cell crosslinking. Commun Chem 8(1), 191 (2025) 10.1038/s42004-025-01568-1

[76] Nouchikian, L., Fernandez-Martinez, D., Renard, P.-Y., Sabot, C., Dumenil, G., Rey, M., Chamot-Rooke, J.: Do not waste TimeEnsure success in your Cross-Linking mass spectrometry experiments before you begin. Anal Chem 96(6), 2506–2513 (2024) 10.1021/acs.analchem.3c04682

[77] Keich, U., Tamura, K., Noble, W.S.: Averaging strategy to reduce variability in Target-Decoy estimates of false discovery rate. J Proteome Res 18(2), 585–593 (2019) 10.1021/acs.jproteome.8b00802

[78] Kalhor, M., Lapin, J., Picciani, M., Wilhelm, M.: Rescoring peptide spectrum matches: Boosting proteomics performance by integrating peptide property predictors into peptide identification. Mol Cell Proteomics 23(7), 100798 (2024) 10.1016/j.mcpro.2024.100798

[79] Tabata, T., Yoshizawa, A.C., Ogata, K., Chang, C.-H., Araki, N., Sugiyama, N., Ishihama, Y.: UniScore, a unified and universal measure for peptide identification by multiple search engines. Mol Cell Proteomics 24(7), 101010 (2025) 10.1016/j.mcpro.2025.101010

[80] Fondrie, W.E., Noble, W.S.: mokapot: Fast and flexible semisupervised learning for peptide detection. J Proteome Res 20(4), 1966–1971 (2021) 10.1021/acs.jproteome.0c01010

[81] Käll, L., Canterbury, J.D., Weston, J., Noble, W.S., MacCoss, M.J.: Semi-supervised learning for peptide identification from shotgun proteomics datasets. Nat Methods 4(11), 923–925 (2007) 10.1038/nmeth1113

[82] Zhao, R., Shen, J., Green, M.R., MacMorris, M., Blumenthal, T.: Crystal structure of UAP56, a DExD/H-box protein involved in pre-mRNA splicing and mRNA export. Structure 12(8), 1373–1381 (2004) 10.1016/j.str.2004.06.006

[83] Henn, A., Bradley, M.J., De La Cruz, E.M.: ATP utilization and RNA conformational rearrangement by DEAD-box proteins. Annu Rev Biophys 41, 247–267 (2012) 10.1146/annurev-biophys-050511-102243

[84] Yellamaty, R., Sharma, S.: Critical cellular functions and mechanisms of action of the RNA helicase UAP56. J Mol Biol 436(12), 168604 (2024) 10.1016/j.jmb.2024.168604

[85] Abramson, J., Adler, J., Dunger, J., et al.: Accurate structure prediction of biomolecular interactions with AlphaFold 3. Nature 630, 493–500 (2024) 10.1038/s41586-024-07487-w

[86] Sinnott, M., Malhotra, S., Madhusudhan, M.S., Thalassinos, K., Topf, M.: Combining information from crosslinks and monolinks in the modeling of protein structures. Structure 28(9), 1061–10703 (2020) 10.1016/j.str.2020.05.012

[87] Keller, A., Tang, X., Bruce, J.E.: Integrated analysis of Cross-Links and Dead-End peptides for enhanced interpretation of quantitative XL-MS. J Proteome Res 22(9), 2900–2908 (2023) 10.1021/acs.jproteome.3c00191

[88] Giese, S.H., Sinn, L.R., Wegner, F., Rappsilber, J.: Retention time prediction using neural networks increases identifications in crosslinking mass spectrometry. Nat Commun 12(1), 3237 (2021) 10.1038/s41467-021-23441-0

[89] Kalhor, M., Saylan, C.C., Picciani, M., Fischer, L., Schimweg, F.B., Lapin, J., Rappsilber, J., Wilhelm, M.: Prosit-XL: enhanced cross-linked peptide identification by fragment intensity prediction to study protein interactions and structures. Nat Commun 16(1), 5429 (2025) 10.1038/s41467-025-61203-4

[90] Chen, M., Hao, Y., Huang, X., Wu, P., Sun, J., Zhang, B., Chen, S.: XL-MSDigger: a deep learning-based, versatile solution for cross-linking mass spectrometry. Nat Commun 17(1) (2026) 10.1038/s41467-026-69489-8

[91] Deng, W., Shi, X., Tjian, R., Lionnet, T., Singer, R.H.: CASFISH: CRISPR/Cas9-mediated in situ labeling of genomic loci in fixed cells. Proc. Natl. Acad. Sci. U. S. A. 112(38), 11870–11875 (2015) 10.1073/pnas.1515692112

[92] Rappsilber, J., Mann, M., Ishihama, Y.: Protocol for micro-purification, enrichment, pre-fractionation and storage of peptides for proteomics using StageTips. Nature Protocols 2(8), 1896–1906 (2007) 10.1038/nprot.2007.261

[93] Velez-Aguilera, G., Ossareh-Nazari, B., Pintard, L.: Dissecting the multiple functions of the Polo-Like kinase 1 in the c. elegans zygote. Cell Cycle Control, 63–88 (2024) 10.1007/978-1-0716-3557-54

[94] Silva, N., Ferrandiz, N., Barroso, C., Tognetti, S., Lightfoot, J., Telecan, O., Encheva, V., Faull, P., Hanni, S., Furger, A., Snijders, A., Speck, C., Martinez-Perez, E.: The fidelity of synaptonemal complex assembly is regulated by a signaling mechanism that controls early meiotic progression. Developmental Cell 31(4), 503–511 (2014) 10.1016/j.devcel.2014.10.001

[95] Wisniewski, J.R., Zougman, A., Nagaraj, N., Mann, M.: Universal sample preparation method for proteome analysis. Nat. Methods 6(5), 359–362 (2009) 10.1038/nmeth.1322

[96] Dorfer, V., Pichler, P., Stranzl, T., Stadlmann, J., Taus, T., Winkler, S., Mechtler, K.: Ms amanda, a universal identification algorithm optimized for high accuracy tandem mass spectra. Journal of Proteome Research 13(8), 3679–3684 (2014) 10.1021/pr500202e

[97] Pirklbauer, G.J., Stieger, C.E., Matzinger, M., Winkler, S., Mechtler, K., Dorfer, V.: MS annika: A new Cross-Linking search engine. J. Proteome Res. (2021) 10.1021/acs.jproteome.0c01000

[98] Birklbauer, M.J., Matzinger, M., Müller, F., Mechtler, K., Dorfer, V.: MS annika 2.0 identifies Cross-Linked peptides in MS2–MS3-Based workflows at high sensitivity and specificity. J. Proteome Res. (2023) 10.1021/acs.jproteome.3c00325

[99] Mendes, M.L., Fischer, L., Chen, Z.A., Barbon, M., O’Reilly, F.J., Giese, S.H., Bohlke-Schneider, M., Belsom, A., Dau, T., Combe, C.W., Graham, M., Eisele, M.R., Baumeister, W., Speck, C., Rappsilber, J.: An integrated workflow for crosslinking mass spectrometry. Mol Syst Biol 15(9), 8994 (2019) 10.15252/msb.20198994

[100] Pettersen, E.F., Goddard, T.D., Huang, C.C., Meng, E.C., Couch, G.S., Croll, T.I., Morris, J.H., Ferrin, T.E.: Ucsf chimerax: Structure visualization for researchers, educators, and developers. Protein Science 30(1), 70–82 (2021) 10.1002/pro.3943. Epub 2020 Oct 22

[101] Lagerwaard, I.M., Albanese, P., Jankevics, A., Scheltema, R.A.: Xlink mapping and analysis (xmas) −smooth integrative modeling in chimerax. bioRxiv (2022) 10.1101/2022.04.21.489026 https://www.biorxiv.org/content/early/2022/06/29/2022.04.21.489026.full.pdf

[102] Birklbauer, M.J., Buur, L.M., Kaser, S., Müller, F., Matzinger, M., Mechtler, K., Winkler, S., Dorfer, V.: Unified down-stream analysis of crosslinking mass spectrometry results with pyXLMS. bioRxiv (2025) 10.64898/2025.12.18.695169

[103] Nelli, F.: Pandas in 7 Days: Utilize Python to Manipulate Data, Conduct Scientific Computing, Time Series Analysis, and Exploratory Data Analysis. BPB Publications (2022)

[104] McKinney, W.: Data structures for statistical computing in python. SciPy 2010 (2010) 10.25080/Majora-92bf1922-00a

[105] Harris, C.R., Millman, K.J., Walt, S.J., Gommers, R., Virtanen, P., Cournapeau, D., Wieser, E., Taylor, J., Berg, S., Smith, N.J., Kern, R., Picus, M., Hoyer, S., Kerkwijk, M.H., Brett, M., Haldane, A., Del Río, J.F., Wiebe, M., Peterson, P., Gerard-Marchant, P., Sheppard, K., Reddy, T., Weckesser, W., Abbasi, H., Gohlke, C., Oliphant, T.E.: Array programming with NumPy. Nature 585(7825), 357–362 (2020) 10.1038/s41586-020-2649-2

[106] Hunter, J.D.: Matplotlib: A 2D graphics environment. Comput. Sci. Eng. 9(3), 90–95 (2007) 10.1109/mcse.2007.55

[107] Waskom, M.: seaborn: statistical data visualization. J. Open Source Softw. 6(60), 3021 (2021) 10.21105/joss.03021

[108] Garreta, R., Moncecchi, G.: Learning Scikit-Learn: Machine Learning in Python. Packt Pub Limited,(2013)

[109] Virtanen, P., Gommers, R., Oliphant, T.E., Haberland, M., Reddy, T., Cournapeau, D., Burovski, E., Peterson, P., Weckesser, W., Bright, J., Walt, S.J., Brett, M., Wilson, J., Millman, K.J., Mayorov, N., Nelson, A.R.J., Jones, E., Kern, R., Larson, E., Carey, C.J., Polat, İ., Feng, Y., Moore, E.W., VanderPlas, J., Laxalde, D., Perktold, J., Cimrman, R., Henriksen, I., Quintero, E.A., Harris, C.R., Archibald, A.M., Ribeiro, A.H., Pedregosa, F., Mulbregt, P., SciPy 1.0 Contributors: SciPy 1.0: fundamental algorithms for scientific computing in python. Nat. Methods 17(3), 261–272 (2020) 10.1038/s41592-019-0686-2

[110] Cock, P.J.A., Antao, T., Chang, J.T., Chapman, B.A., Cox, C.J., Dalke, A., Friedberg, I., Hamelryck, T., Kauff, F., Wilczynski, B., Hoon, M.J.L.: Biopython: freely available python tools for computational molecular biology and bioinformatics. Bioinformatics 25(11), 1422–1423 (2009) 10.1093/bioinformatics/btp163

[111] Hulstaert, N., Shofstahl, J., Sachsenberg, T., Walzer, M., Barsnes, H., Martens, L., Perez-Riverol, Y.: ThermoRawFileParser: Modular, scalable, and Cross-Platform RAW file conversion. J Proteome Res 19(1), 537–542 (2020) 10.1021/acs.jproteome.9b00328

[112] Goloborodko, A.A., Levitsky, L.I., Ivanov, M.V., Gorshkov, M.V.: Pyteomics–a python framework for exploratory data analysis and rapid software prototyping in proteomics. J Am Soc Mass Spectrom 24(2), 301–304 (2013) 10.1007/s13361-012-0516-6

[113] Levitsky, L.I., Klein, J.A., Ivanov, M.V., Gorshkov, M.V.: Pyteomics 4.0: Five years of development of a python proteomics framework. J Proteome Res 18(2), 709–714 (2019) 10.1021/acs.jproteome.8b00717

[114] Schilling, B., Rardin, M.J., MacLean, B.X., Zawadzka, A.M., Frewen, B.E., Cusack, M.P., Sorensen, D.J., Bereman, M.S., Jing, E., Wu, C.C., Verdin, E., Kahn, C.R., Maccoss, M.J., Gibson, B.W.: Platform-independent and label-free quantitation of proteomic data using MS1 extracted ion chromatograms in skyline: application to protein acetylation and phosphorylation: Application to protein acetylation and phosphorylation. Mol. Cell. Proteomics 11(5), 202–214 (2012) 10.1074/mcp.M112.017707

[115] Perez-Riverol, Y., Bai, J., Bandla, C., García-Seisdedos, D., Hewapathirana, S., Kamatchinathan, S., Kundu, D.J., Prakash, A., Frericks-Zipper, A., Eisenacher, M., Walzer, M., Wang, S., Brazma, A., Vizcaíno, J.A.: The PRIDE database resources in 2022: a hub for mass spectrometry-based proteomics evidences. Nucleic Acids Res 50(D1), 543–552 (2022) 10.1093/nar/gkab1038

[116] Perez-Riverol, Y., Bandla, C., Kundu, D.J., Kamatchinathan, S., Bai, J., Hewapathirana, S., John, N.S., Prakash, A., Walzer, M., Wang, S., Vizcaíno, J.A.: The PRIDE database at 20 years: 2025 update. Nucleic Acids Res (2024) 10.1093/nar/gkae1011

[117] Combe, C.W., Graham, M., Kolbowski, L., Fischer, L., Rappsilber, J.: xiview: Visualisation of crosslinking mass spectrometry data. Journal of Molecular Biology 436(17), 168656 (2024) 10.1016/j.jmb.2024.168656. Computation Resources for Molecular Biology

